# Heterogeneity in surface sensing produces a division of labor in *Pseudomonas aeruginosa* populations

**DOI:** 10.1101/532598

**Authors:** Catherine R. Armbruster, Calvin K. Lee, Jessica Parker-Gilham, Jaime de Anda, Aiguo Xia, Boo Shan Tseng, Lucas R. Hoffman, Fan Jin, Caroline S. Harwood, Gerard C. L. Wong, Matthew R. Parsek

## Abstract

The second messenger signaling molecule cyclic diguanylate monophosphate (c-di-GMP) drives the transition from planktonic to biofilm growth in many bacterial species. *Pseudomonas aeruginosa* has two surface sensing systems that produce c-di-GMP in response to surface adherence. The current thinking in the field is that once cells attach to a surface, they uniformly respond with elevated c-di-GMP. Here, we describe how the Wsp system generates heterogeneity in surface sensing, resulting in two physiologically distinct subpopulations of cells. One subpopulation has elevated c-di-GMP and produces biofilm matrix, serving as the founders of initial microcolonies. The other subpopulation has low c-di-GMP and engages in surface motility, allowing for exploration of the surface. We also show that this heterogeneity strongly correlates to surface behavior for descendent cells. Together, our results suggest that after surface attachment, *P. aeruginosa* engages in a division of labor that persists across generations, accelerating early biofilm formation and surface exploration.

## Introduction

*Pseudomonas aeruginosa* is an opportunistic pathogen that engages in a range of surface-associated behaviors and is a model bacterium for studies of surface-associated communities called biofilms. Biofilms are dense aggregates of cells producing extracellular matrix components that hold the community together. The biofilm mode of growth confers cells protection from a variety of environmental stresses including nutrient limitation, desiccation, and shear forces, as well as engulfment by protozoa in the environment or phagocytes in a host (*1*).

The secondary messenger signaling molecule cylic-di-GMP (c-di-GMP) drives the transition from the planktonic to the biofilm mode of growth. In many bacterial species, including *P. aeruginosa*, elevated c-di-GMP results in repression of flagellar motility genes, while promoting expression of genes involved in producing a biofilm matrix(*2*). The *P. aeruginosa* biofilm matrix is composed of a combination of polysaccharides (including Pel and Psl), proteins (including the adhesin CdrA), and extracellular DNA (*3*–*8*). Biofilm matrix production is an energetically costly process that is regulated at multiple levels (*9*). The *cdrA, pel* and *psl* genes are all transcriptionally induced under conditions of high c-di-GMP(*10*).

For many species, the initial step in biofilm formation involves adherence of free swimming planktonic cells to a surface and the initiation of surface sensing. *P. aeruginosa* has at least two distinct surface sensing systems, the Wsp and the Pil-Chp systems, that when activated, lead to biofilm formation. The Wsp system senses an unknown surface-related signal (recently proposed to be membrane perturbation (*11*)) through WspA, a membrane-bound protein homologous to methyl-accepting chemotaxis proteins (MCPs). Activation of this system stimulates phosphorylation of the diguanylate cyclase WspR, which leads to the formation of aggregates of phosphorylated WspR (WspR-P) in the form of visible subcellular clusters. This aggregation of WspR-P potentiates its activity, increasing c-di-GMP synthesis (*12*). In comparison, the Pil-Chp chemosensory-like system initiates a hierarchical cascade of second messenger signaling in response to a surface (*13*). First, an increase in cellular cAMP levels occurs through activation of the adenylate cyclase CyaB by the chemotaxis-like Pil-Chp complex. This increases expression of genes involved in type IV pilus biogenesis, including PilY1. PilY1 is associated with the type IV pilus and harbors a Von Willebrand motif, which is involved in mechanosensing in eukaryotic systems(*14*). Thus, it has been proposed that this protein may be involved in the mechanosensing of surfaces (*15*). The output of this second signal is through the diguanylate cyclase, SadC, resulting in an increase in cellular c-di-GMP levels. Unlike the Wsp system, which localizes laterally along the cell (*16*), PilY1 is required to be associated with polarly-localized type IV pili in order to stimulate c-di-GMP production (*13*, *14*), suggesting that *P. aeruginosa* deploys both polar and laterally localized systems to promote c-di-GMP synthesis in response to a surface.

Here, we examined the dynamics of c-di-GMP production and bacterial surface motility at the single-cell level during early stages of biofilm formation. We used a plasmid-based, transcriptional reporter of intracellular c-di-GMP to follow the downstream fate of cells producing varying levels of c-di-GMP in response to surface attachment. Within a clonal population of *P. aeruginosa*, we found that levels of c-di-GMP vary among individual cells as they sense a surface, leading to a division of labor between two energetically costly behaviors associated with early biofilm formation: surface exploration and polysaccharide production.

## Results

### Cellular c-di-GMP levels rapidly increase upon surface attachment

We initially compared levels of c-di-GMP between *P. aeruginosa* PAO1 cells growing attached to a silicone surface and subjected to constant flow for 4 hours to those grown planktonically for 4 h. As expected, we observed that PAO1 cellular c-di-GMP levels are 4.4-fold higher (± 0.78 SD, N = 3, p ≤ 0.05) after 4h of growth attached to a surface compared to planktonic growth (Figure 1A). Because direct measurement of c-di-GMP by LC-MS/MS is limited by our ability to generate enough biomass at earlier time points, we used qRT-PCR to monitor *pel* transcript levels as a readout of c-di-GMP. We found that after just 30 min of surface attachment, *pelA* transcript levels had increased almost 10-fold compared to planktonically grown cells (Figure 1 – Supplement 1). This is consistent with previously published literature showing that transcription of the *pel* operon is directly and positively controlled by high cellular levels of c-di-GMP (*17*, *18*).

**Figure 1.**
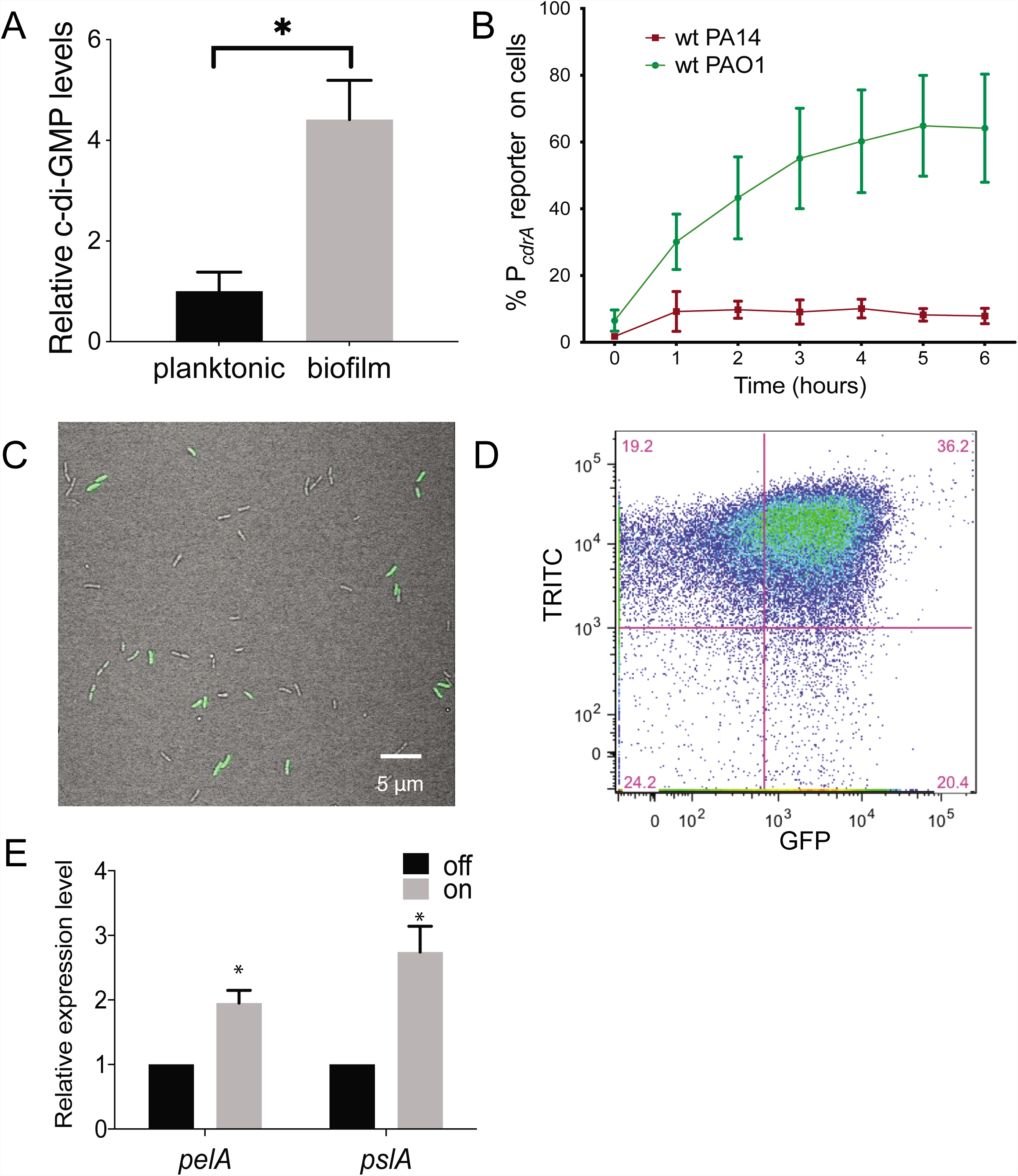
Heterogenetity in cellular levels of c-di-GMP during early *P. aeruginosa* biofilm formation. (A) c-di-GMP levels are elevated rapidly upon association of *P. aeruginosa* PAO1 cells with a surface. Relative levels of intracellular c-di-GMP in wild type PAO1 cells grown either planktonically or after 4 h of attachment to a silicone tube. Values are normalized to the average concentration of c-di-GMP in planktonic cells, in pmol c-di-GMP/mg total protein as determined by LC-MS/MS, and presented as mean and SD. * p < 0.05 by T-test, N= 3. Figure 1 – Figure supplement 1 shows the Pel polysaccharide operon is transcriptionally activated almost 10-fold compared to planktonic cells within 30 minutes of surface attachment. (B) Two commonly studied *P. aeruginosa* lab strains, PAO1 and PA14, differentially activate the c-di-GMP reporter during surface sensing. Wild type PAO1 or PA14 cells harboring the c-di-GMP reporter (P_*cdrA*_::*gfp*_ASV_) were grown to mid-log phase in planktonic culture, then inoculated into a flow cell and supplied with 1% LB medium. Surface attached cells were imaged immediately after inoculation (time 0 h), and hourly for 12 hours. The c-di-GMP reporter is activated in a subset of wild type PAO1 cells within 1 hour of surface attachment and remains activated in approximately 60% of PAO1 cells during the first 6 hours of attachment. In PA14, the c-di-GMP reporter is activated in a smaller proportion of attached cells compared to PAO1. Data points are mean percentage of reporter activated cells from each time point across at least 3 biological replicates, with standard deviation. Figure 1 – Supplement 4 shows an additional c-di-GMP responsive transcriptional reporter (using the *siaA* promoter) is also responsive to Wsp-dependent changes in cellular levels of c-di-GMP. (C) Wild type PAO1 cells display heterogeneity in c-di-GMP reporter activity after 6 hours of surface attachment. Confocal microscopy image of wild type PAO1 P_*cdrA*_::*gfp*_ASV_ grown in 1% LB after 6 hours of surface attachment during a time course flow cell experiment. Bright field (grey) and GFP (green) channels are merged. Wild type PAO1 P_*cdrA*_::*gfp*_ASV_ was grown in 1% LB and imaged by CSLM. Figure 1 – Figure supplement 2 shows additional representative timecourse images of PAO1. (D) Psl exopolysaccharide production is enriched in the population of cells with high c- di-GMP. Representative scatterplot of reporter activity versus Psl lectin binding in wild type PAO1 harboring the pP_*cdrA*_::*gfp*_ASV_ reporter grown for four hours in LB before surface attached cells were harvested, stained with the lectin, washed, and counted by flow cytometry. (E) Subpopulations of PAO1 cells with high and low c-di-GMP reporter activity are physiologically distinct. Cells with higher c-di-GMP reporter activity have increased expression of Pel and Psl biosynthetic machinery genes. After 4 hours of attachment to glass, wild type PAO1 cells were separated by flow-assisted cell sorting (FACS) into a population of cells with high (on) and low (off) c-di-GMP reporter activity, then qRT-PCR was performed to quantify expression of Pel and Psl exopolysaccharide biosynthesis genes. Levels of expression of Pel or Psl mRNA were normalized to the off population. * p < 0.05 by T-test, N= 3 biological replicates. Figure 1 – Figure supplement 3 shows controls for validating the protocol to monitor pP_*cdrA*_::*gfp*_ASV_ by flow cytometry. Figure 1 – Supplement 5 shows by flow cytometry that Psl and Pel polysaccharide production is highest in cells with high pP_*cdrA*_::*gfp*_ASV_ reporter activity.

### The P*_cdrA_*::*gfp* reporter detects heterogeneity in c-di-GMP during surface sensing

Next, we sought to visualize early c-di-GMP signaling events at the single cell level. To this end we used a plasmid-based, c-di-GMP responsive transcriptional reporter, pP_*cdrA*_::*gfp*_ASV_ (*19*) in two commonly-studied *P. aeruginosa* strains, PAO1 and PA14. Planktonic cells (a condition where the reporter is inactive due to low c-d-GMP levels) were used to inoculate flow cell chambers. We imaged individual cells of each reporter strain hourly for up to 6 hours after surface attachment (Figure 1B and Figure 1 – Supplement 2). As expected, we saw minimal GFP fluorescence at the 0 h time point (right after surface attachment). However, by 1 h, the reporter was activated in a subset of surface attached cells, as defined by GFP fluorescence greater than twice that of background levels (referred to as reporter “on” subpopulations). Interestingly, between 4 and 6 h post inoculation, we consistently observed that the c-di-GMP reporter was only active in a subset of cells in both strains (Figure 1C). In PA14, the reporter was activated in 10% of the population over 6 h, whereas PAO1 displayed greater reporter activity, with 40-60% of the cells displaying reporter activity through 12 h (Figure 1 – Supplement 2). We confirmed these results using flow cytometry to assess the proportion of attached cells that were fluorescent (Figure 1 – Supplement 3D,E). To be sure that the promoter of *cdrA* is representative of c-di-GMP-regulated gene expression, we replaced P_*cdrA*_ with the promoter of *siaA*, a gene that is also highly expressed under conditions of elevated c-di-GMP (*10*, *20*). We found that pP_*siaA*_::*gfp*_ASV_ reporter activity resembled that of pP_*cdrA*_::*gfp*_ASV_ in response to a surface (Figure 1 – Supplement 4). Thus, reporter activity is indeed linked to cellular levels of c-di-GMP.

### Cyclic di-GMP heterogeneity leads to phenotypic diversification at early stages of biofilm formation

We then wanted to confirm that subpopulations of surface-attached *P. aeruginosa* cells with high and low c-di-GMP reporter activity are truly physiologically distinct from one another. We used TRITC-labeled lectins to stain for two c-di-GMP-induced exopolysaccharides, Psl and Pel (*7*, *21*), the presence of which is indicative of biofilm formation by PAO1 and PA14, respectively. After 4h of attachment to glass, we observed an enrichment of TRITC-conjugated lectin staining in the population of cells with high c-di-GMP reporter activity (Figure 1D and Figure 1 – Supplement 5), demonstrating that the subpopulation of cells with high c-di-GMP is producing more exopolysaccharide than their low c-di-GMP counterparts. As a complementary approach, we separated 4 h surface-grown cells of the reporter strain into reporter “on” and “off” subpopulations using flow-assisted cell sorting (FACS; Figure 1 – Supplement 6). We then applied qRT-PCR to compare Pel and Psl transcript levels in these two populations. Both the *pel* and *psl* operon transcripts were elevated in the reporter “on” subpopulation, relative to the reporter “off” subpopulation (Figure 1E). These data support that, with respect to c-di-GMP signaling, there are at least two distinct subpopulations that arise shortly after surface attachment.

### The Wsp system is required for surface sensing

We next evaluated the relative contributions of the Wsp and Pil-Chp surface sensing systems to surface-induced c-di-GMP production. Strains with mutations in the Pil-Chp chemosensory system were not significantly defective in surface sensing activity. Deletion of the diguanylate cyclase activated through the Pil-Chp system (PAO1 Δ*sadC*) and the gene encoding the putative sensor PilY1 (PAO1 Δ*pilY1*) did not significantly influence reporter activity in response to a surface (Figure 2 – Supplement 1A,B). Whereas both the SadC and PilY1 mutants displayed wild type levels of reporter activity, a mutant lacking the main Type IV pilus filament protein (PAO1 Δ*pilA*) did show a statistically significant defect in reporter activity by 6 h (Figure 2 – Supplement 1B; p <0.05 by T-test). We then mutated the c-di-GMP cyclase gene, *wspR,* to inactivate the Wsp system. In addition, we deleted the gene encoding the methylesterase *wspF*, which locks the system into the active state, regardless of whether cells are surface-associated. We found that PAO1 Δ*wspR* strain exhibited extremely low levels of reporter activity during the first 6 h after surface attachment (Figure 2A and Figure 2 – Supplement 2). Complementation of PAO1 Δ*wspR* restored wild type levels of activity at all time points (Figure 2 – Supplement 3). As expected, PAO1 Δ*wspF* had a high proportion of reporter active cells (Figure 2A). We repeated these experiments in the lab strain PA14 and saw a similar trend for Wsp and Pil-Chp mutants (Figure 2 – Supplement 4).

**Figure 2.**
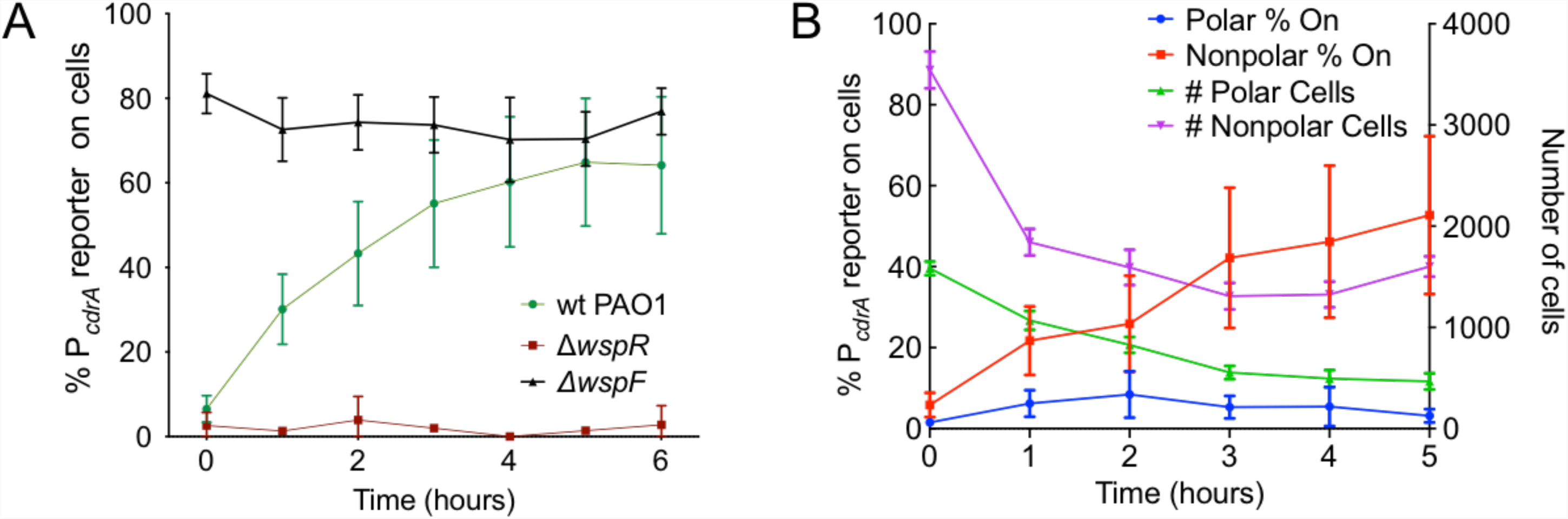
The Wsp system generates heterogeneity in cellular levels of c-di-GMP during early *P. aeruginosa* biofilm formation. (A) The Wsp system is required for activation of the pP_*cdrA*_::*gfp*_ASV_ reporter during surface sensing. Six hour time course plot of the average percentage of surface-attached cells from either wild type PAO1 (green), PAO1 Δ*wspR* (red), or PAO1 Δ*wspF* (black) in which the pP_*cdrA*_::*gfp*_ASV_ reporter had turned “on” at each hour. Cells were identified as “on” if their average GFP fluorescence was greater than twice the average background GFP fluorescence of the image. Error bars = standard deviation. N ≥ 3 biological replicates. See Figure 2 – Figure supplement 1 for the same timecourse using mutants in the Pil-Chp surface sensing system. Figure 2 – Figure supplement 2 shows representative images from Figure 2A. Figure 2 – Figure supplement 3 shows that complementing the *wspR* mutant restores wild type levels of reporter activity. Figure 2 – Figure supplement 4 shows that the lab strain PA14 also displays Wsp-dependent c-di-GMP heterogeneity. (B) Laterally attached cells have higher c-di-GMP levels than polarly attached cells. Five hour time course plot depicting, on the left axis, the percentage of pP_*cdrA*_:*gfp*_ASV_ reporter “on” cells that were either polarly (blue) or laterally (red) attached to the surface of a glass coverslip in a flow cell at each hour. The right axis depicts the total number of polar (green) and laterally attached (purple) cells at each time point. Cells were identified as pP_*cdrA*_:*gfp*_ASV_ reporter “on” if their average GFP fluorescence was greater than twice the average background GFP fluorescence of the image. Error bars = standard deviation. N = 4 biological replicates.

Since the Pil-Chp surface sensing apparatus is polarly localized and the Wsp system is localized laterally along the length of the cell body, we examined whether reporter activity correlated with polar versus lateral attachment to the surface. We found that reporter activity was very low in polarly attached cells, while cells attached along the entire length of the cell body displayed a higher proportion of activated cells (Figure 2B). This finding is also consistent with the localization of the Wsp system and its role for early c-di-GMP signaling during surface sensing.

### Heterogeneity in c-di-GMP levels among cells correlates with Wsp system activity

The specific activity of purified WspR increases as a function of WspR concentration when the protein is treated with beryllium fluoride to mimic phosphorylation, supporting the idea that formation of subcellular clusters of WspR-P potentiates its diguanylate cyclase activity and leads to elevated c-di-GMP(*12*). Fewer than 1% of wild-type cells grown in broth have a visible WspR-YFP cluster. However, after a short period of growth on an agar surface, WspR-YFP clusters were visible in 30-40% of wild type PAO1 cells, and this is dependent on sensing by the membrane-bound protein WspA, which is laterally distributed in cells (9). To directly link WspR cluster formation with diguanylate cyclase activity at the cellular level and with surface sensing, we constructed a version of the c-di-GMP reporter that expresses mTFP1 instead of GFP (pP_*cdrA*_::*mTFP1*) to avoid the issue of spectral overlap with WspR-YFP. We monitored reporter activity in two point mutants of WspR (L170D and E253A) that are driven by an inducible promoter, translationally fused to eYFP and have been previously shown to form large subcellular WspR clusters in a higher percentage of cells than wild-type WspR. The WspR[L170D] protein is highly active for c-di-GMP production, and it forms subcellular clusters in about 75% of agar surface-grown cells. A WspR[E253A] point mutation abolishes diguanylate cyclase activity, but this protein still forms clusters in about 70% of surface-grown cells (*12*). As expected, in the presence of inducer, we observed a large increase in c-di-GMP reporter activity in WspR[L170D], but not WspR[E253A] (Figure 3A, B). We then asked whether the heterogeneity in reporter activity in response to surface attachment correlates with WspR clustering in the WspR[L170D] strain. We found that pP_*cdrA*_::*mTFP1* activity was significantly higher in cells with at least one subcellular WspR-eYFP focus in the WspR[L170D] strain compared to cells without a WspR-eYFP focus (Figure 3C; median mTFP1 fluorescence of 345 vs. 320 RFU respectively, Mann-Whitney test, p < 0.001). These data indicate that the heterogeneity observed in c-di-GMP signaling after surface attachment is due to the heterogeneity in the activity of the Wsp system, as reflected by subcellular clustering of active WspR-P.

**Figure 3.**
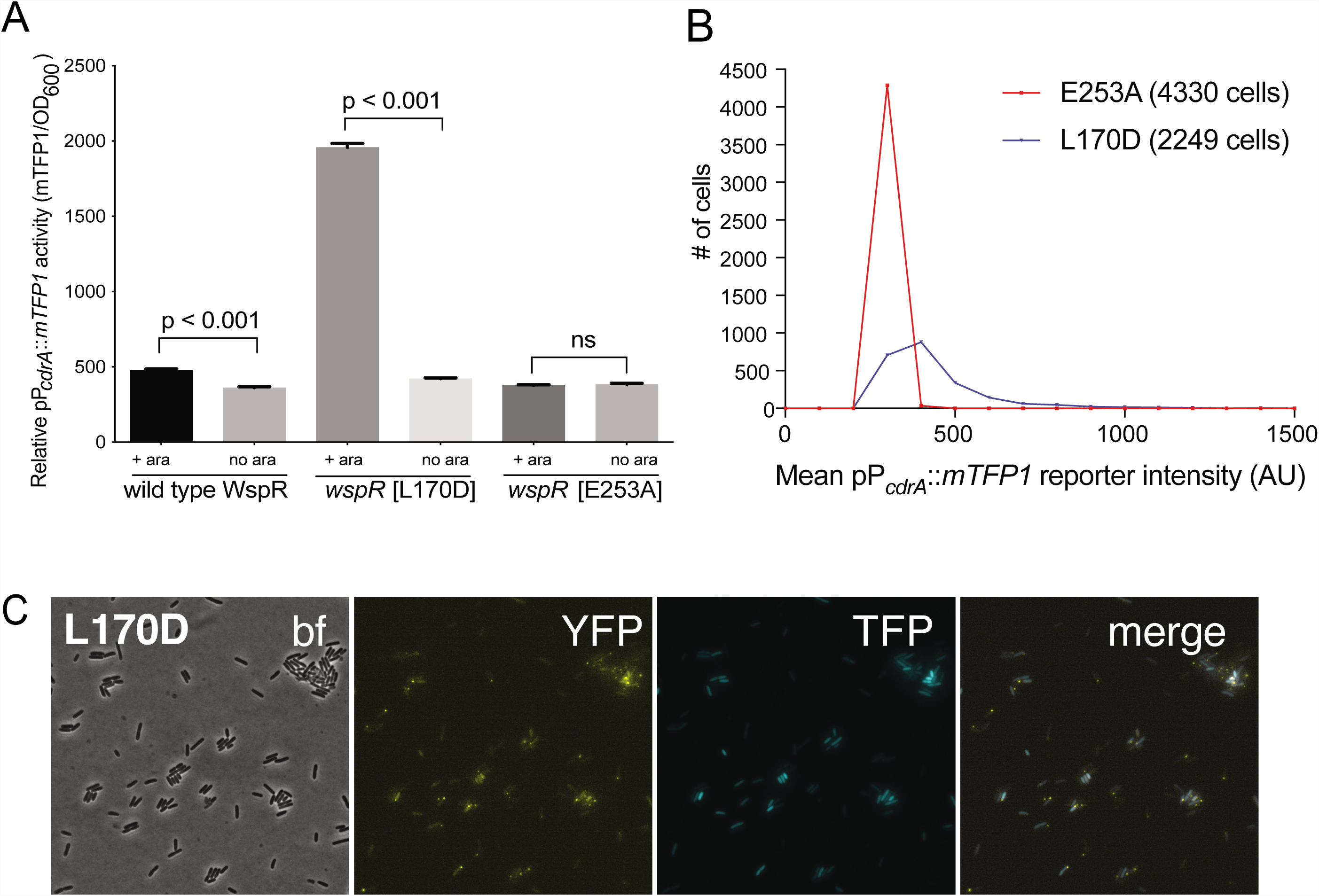
Activity of the pP***cdrA***::*mTFP1* reporter is dependent on the ability of WspR to produce c-di-GMP. (A) The pP_*cdrA*_::*mTFP1* reporter is active in surface grown cells with functional, arabinose-inducible alleles of WspR when arabinose is added to the media. Wild type WspR represents the strain PAO1 Δ*wspR* attCTX::*wspR*-eYFP. *wspR*[L170D] represents the strain PAO1 Δ*wspR* attCTX::*wspR*[L170D]-eYFP, which produces large subcellular clusters of WspR and grows as rugose small colonies on LB with 1% arabinose, a phenotype that is indicative of high intracellular c-di-GMP. *wspR*[E253A] represents the strain PAO1 Δ*wspR* attCTX::*wspR*[L170D]-eYFP cells, which forms large subcellular WspR clusters, but does not produce c-di-GMP via WspR due to the point mutation located in its active site. Cells were grown on LB agar plates with 100 µg/mL gentamicin, and in the presence or absence of 1% arabinose. Cells were resuspended in PBS and mTFP1 fluorescence and OD_600_ were measured. Relative pP_*cdrA*_::*mTFP1* reporter activity is the level of mTFP1 fluorescence normalized to OD_600_. Asterisk indicates statistical significance by Student’s t-test (p < 0.001) in 6 technical replicates. Error bars = standard deviation. (B) The pP_*cdrA*_::*mTFP1* reporter displays heterogeneity in a strain with a functional WspR (*wspR*[L170D]) and is consistently dark in a strain with inactive WspR. Histogram displaying the distribution of average cellular levels of mTFP1 fluorescence from expression of the pP_*cdrA*_::*mTFP1* reporter in either the PAO1 Δ*wspR* attCTX::*wspR*[L170D]-eYFP (blue) or PAO1 Δ*wspR* attCTX::*wspR*[L170D]-eYFP (red) backgrounds. (C) Cells with visible subcellulars clusters of WspR[L170D]-eYFP (strain PAO1 Δ*wspR* attCTX::*wspR*[L170D]-eYFP) also have high levels of c-di-GMP reporter activity. bf, bright field; YFP, wspR-YFP foci; mTFP1 = pP_*cdrA*_::*mTFP1* activity; and merge, merged YFP and TFP channels. PAO1 Δ*wspR* attCTX::*wspR*[L170D]-eYFP cells harboring the pP_*cdrA*_::mTFP1 reporter were grown on LB agar plates with 1% arabinose and 100 µg/mL gentamicin, then spotted onto an agar pad and imaged immediately.

We next asked whether the observed heterogeneity in c-di-GMP signaling in response to a surface has a meaningful influence on biofilm formation. This was particularly important since previous published results indicated that a *wspR* mutation had only a small impact on biofilm production (*22*). However, these studies assessed biofilm formation at later stages of biofilm growth that were well beyond initial surface attachment. Therefore, we chose to compare a *wspR* mutant to wild type at earlier biofilm stages. We performed *in vitro* biofilm assays and observed that a PAO1 Δ*wspR* mutant was defective for biofilm formation relative to wild type PAO1 at 2, 4, and 6 hours post-attachment (Figure 4A). However, at later stages of development (∼24 h), the *wspR* mutant caught up and produced similar amounts of biofilm biomass relative to wild type levels. Complementation of the Δ*wspR* strain *in trans* restored wild type levels of biofilm formation at all time points. These data suggest that the Wsp system rapidly responds to surface contact to generate elevated levels of c-di-GMP, which accelerates biofilm production. Given the importance of c-di-GMP signaling in biofilm production, the fact that the Δ*wspR* strain can ultimately attain wild-stype levels of biofilm biomass suggests that one of the many other known c-di-GMP cyclases present in *P. aeruginosa* may ultimately compensate for c-di-GMP production in the absence of WspR.

**Figure 4.**
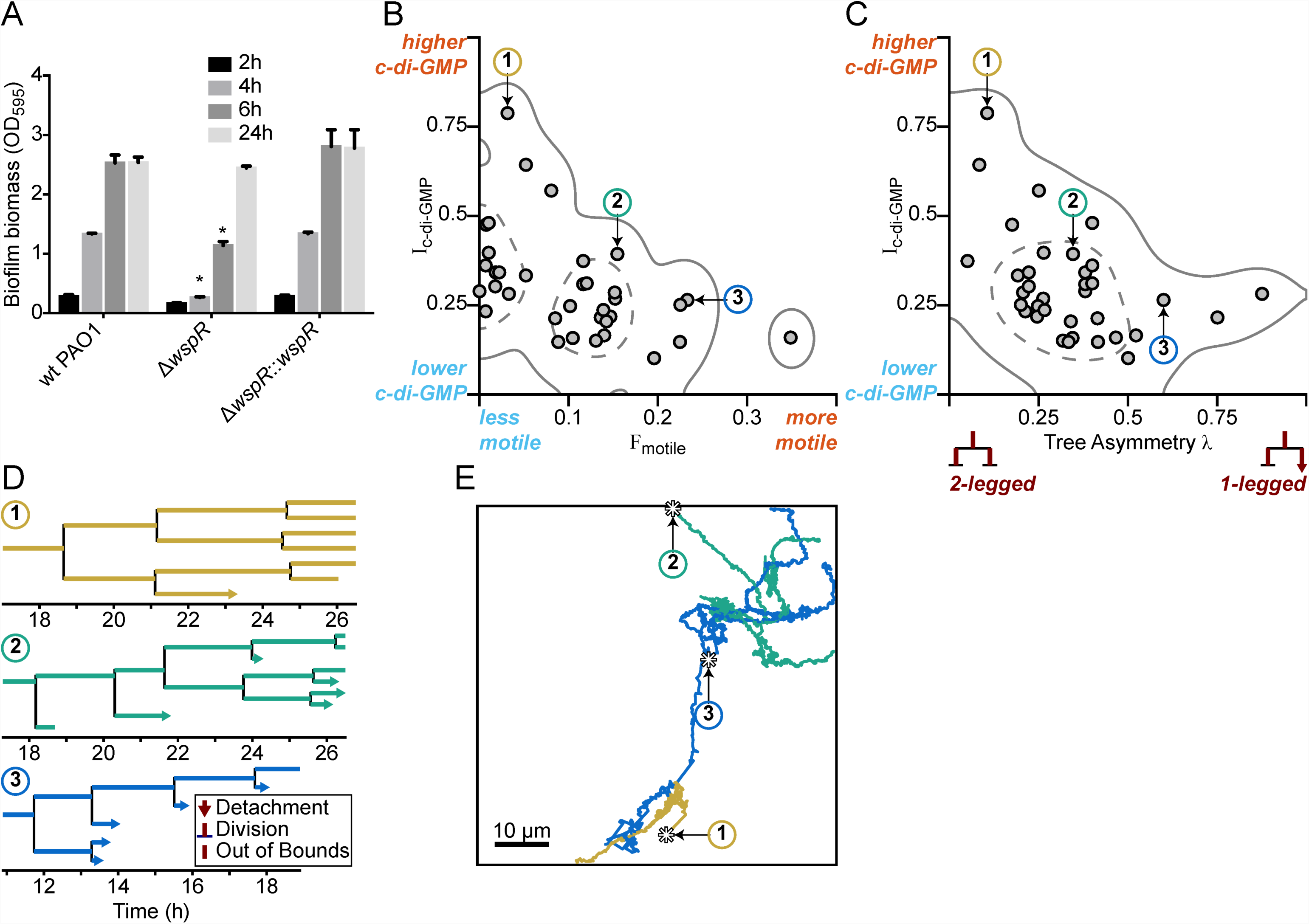
Multigenerational c-di-GMP levels within populations of surface-attached wild type PAO1 cells are inversely correlated with surface motility and detachment. (A) The Wsp surface sensing system is involved in the early stages of biofilm formation in PAO1. Static biofilm assay performed in wild type PAO1, a single deletion mutant of *wspR*, and the PAO1 Δ*wspR* mutant complemented with w*spR*. Between 4 and 6 hours, PAO1 Δ*wspR* shows a defect in surface attachment and biofilm formation relative to the wild type. However, after 24 hours, PAO1 Δ*wspR* formed equal biofilm biomass compared to wild type. Plotted values are the mean of 6 technical replicates and error is standard deviation. Asterisk indicates a statistically significant change in biomass relative to wild type PAO1 at each time point (Student’s t test; p < 0.05). (B) Plot of I_c-di-GMP_ vs F_motile_ for individual wild type PAO1 families. I_c-di-GMP_ is the relative normalized c-di-GMP reporter intensity averaged across all members of a family. F_motile_ is the fraction of time that cells in a family are motile (specifically surface translational motility). Each circle represents an individual family (N = 35) with at least 4 tracked generations. Solid lines represent the 95% probability bounds and dashed lines represent the 50% probability bounds, calculated via kernel density estimation. Spearman correlation: ρ = −0.53, p = 0.0012. (C) Plot of I_c-di-GMP_ vs tree asymmetry λ for individual wild type PAO1 families. Colored numbers indicate the same 3 families from (B) and (D). Tree asymmetry λ quantifies the detachment behavior of family trees as follows. λ = 0 corresponds to ideal trees with purely “two-legged” division-branching, when both daughter cells remain attached to the surface. λ = 1 corresponds to ideal trees with purely “one-legged” division-branching when one daughter cell detaches or travels outside the field of view. Points here are the same families as in (B). Solid lines represent the 95% probability bounds and dashed lines represent the 50% probability bounds, calculated via kernel density estimation. Spearman correlation: ρ = −0.45, p = 0.0068. (D) Family trees of the same 3 representative wild type PAO1 families indicated in (B) and (C). Time 0 h is the start of the dataset recording. Lengths of horizontal lines on the plots are proportional to time spent in each generation. Horizontal lines that end with arrows are detachment events, lines that intersect with a vertical line are division events, and lines that end without a marker are out-of-bound events where we lose track of the bacterium (moving out of the field of view or reaching the end of the recording; represented as moving outside the XYT limits of the dataset boundaries). Vertical lines are arbitrarily spaced to show all the descendants. Colors represent the families in (B) and (C). (E) Spatial trajectories of the 3 representative families. Asterisks (*) represent the initial location of the founder cell. Scale bar 10 µm. The families are color coded as in the previous panels.

### Cyclic di-GMP heterogeneity leads to diversification in surface exploration at the lineage level

We hypothesized that heterogeneity in c-di-GMP signaling dictated by the Wsp complex could impact the surface behavior of the two observed subpopulations. We predicted that the subpopulation of cells with high c-di-GMP after surface attachment would produce biofilm matrix exopolysaccharides and contribute to initial microcolony formation, while the cells with low c-di-GMP would exhibit increased surface motility and detachment, which is known to be inhibited by exopolysaccharide production. To test this hypothesis, we tracked both reporter activity and surface behavior for cells within a single field of view for 40 h. From our single-cell tracking data, we generated family trees across at least four generations of cells, using a previously described technique (*23*). We tracked the time-averaged P_*cdrA*_::*gfp*_ASV_ reporter activity (I_c-di-GMP_), surface motility behavior (F_motile_, defined as the fraction of time that cells are motile), and detachment behavior (tree asymmetry λ where λ = 0 represents both daughter cells remaining attached to the surface and λ = 1 represents when one daughter cell detaches or travels outside the field of view).

In *P. aeruginosa*, surface exploration is mainly accomplished by twitching motility, mediated by type IV pili, and does not appear to be influenced by levels of intracellular c-di-GMP when analyzing single cells (*24*). Interestingly, we found that correlations between c-di-GMP and motility during the lifetime of individual cells are weak. However, when analyzing entire lineages in family trees rather than individual cells, we found clear inverse correlations between I_c-di-GMP_ and F_motile_ (Figure 4B, ρ = - 0.53, p = 0.0012) and between I_c-di-GMP_ and λ (Figure 4C, ρ = −0.45, p = 0.0068), suggesting that c-di-GMP levels is strongly inversely correlated with surface motility behavior and detachment behavior over multiple generation of cells. To illustrate these correlations, we chose three representative families, with either high, intermediate, or low I_c-di-GMP_ and plotted their family trees (Figure 4D) and spatial trajectories (Figure 4E). Families with the highest I_c-di-GMP_ had the lowest F_motile_ and λ (Family 1, Figure 4B-E). In these families, daughter cells remained attached following cell division, exhibited continuously elevated c-di-GMP, did not move appreciable distances on the surface, and ultimately produced small microcolonies. In contrast, families of cells with low I_c-di-GMP_ had the highest F_motile_ and λ. For these families, daughter cells frequently detached or traveled outside the field of view, had lower c-di-GMP levels, traveled larger distances on the surface, and ultimately did not form microcolonies (Family 3, Figure 4B-E).

One important question is what happens to early biofilm development if we were to effectively remove heterogeneity in c-diGMP output rooted in the WspR surface sensing system. To address this question, we used a strain in which c-di-GMP production could be easily controlled using an optogenetic system. The precise control of c-di-GMP expression in individual cells was made possible by the use of a chimeric protein that fused a diguanylate cyclase domain to a bacteriophytochrome domain. Flow chambers were seeded with the optogenetic strain encoding a heme oxygenase (*bphO*) and light-responsive diguanylate cyclase (*bphS*)(*25*). Initially, cells attached on the glass surface were tracked and continuously stimulated with red-light over ∼8 h using adaptive tracking illumination microscopy (ATIM), which allows for precise stimulation of the initial attached cells and their offspring and ensures sustained intracellular c-di-GMP production for a fixed number of surface cell generations (Figure 5 – Supplement 1). Cellular lineages (a cell and all of its offspring) and c-di-GMP expressions were continually monitored for at least 12 h. Families that were not stimulated with light demonstrated a heterogeneous surface response (Figure 5B,D) similar to that of Families 1-3 in Figure 4B-E. Some lineages were dominated by surface explorers, whereas others were seen to commit to microcolony formation. In contrast, in families stimulated with light for more than 1 generation, the resulting c-di-GMP production artificially forced lineages to have low surface motility and commit to microcolony production (Figure 5A,C) similar to that of Family 1 in Figure 4B-E. Families stimulated with light in this manner had higher I_c-di-GMP_ and lower λ values than those that were not stimulated (Figure 5 – Supplement 2). We also found that optogenetic control of c-di-GMP results in phenotypes that are consistent with the wild-type behavior presented in Figure 4, with illuminated cells (high c-di-GMP) displaying the least motility and control (non-illuminated) displaying comparatively greater surface motility (Figure 5 – Supplement 2). Interestingly, families stimulated with light for 1 generation or less are not significantly different from un-illuminated controls (data not shown). Our data show that the generation of c-di-GMP can deterministically lead to the creation of an entire lineage of sessile cells with post-division surface persistence, low motility, and initiation of microcolony formation. Altogether, these results show that c-di-GMP levels, surface motility, and detachment are inversely correlated at the lineage level, and that the time scale for this occurs over multiple generations.

**Figure 5.**
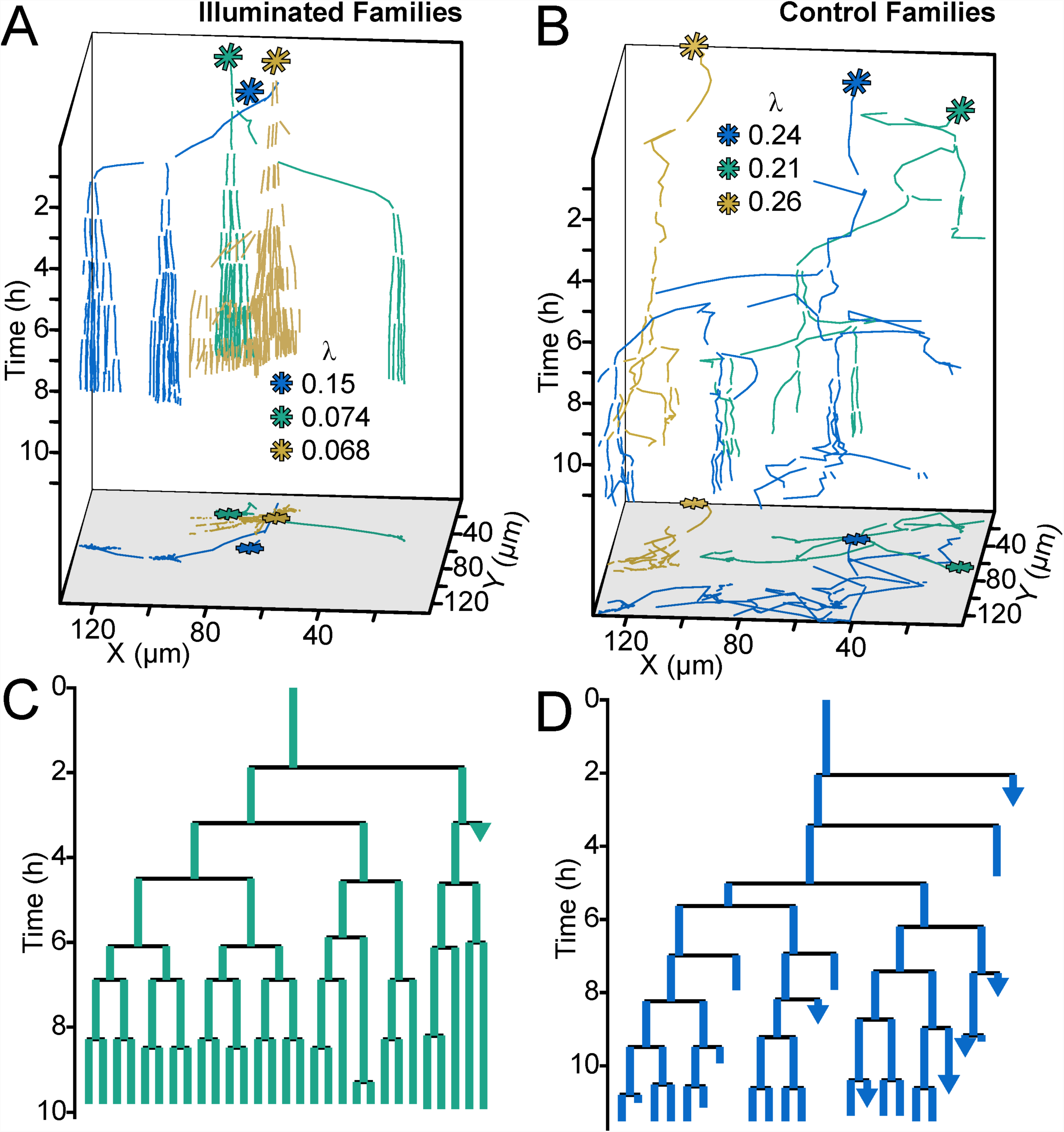
Optogenetic control of c-di-GMP production drastically affects family architecture and surface motility. (A,B) Spatiotemporal plot of 3 illuminated families (A) and 3 control families (B). The individual cell tracks in the 3D plot are projected onto the XY plane as spatial trajectories. As in figure 4, λ is a measure of tree asymmetry, with higher values indicating more cells traveling outside the field of view or detaching. In A, the illuminated families tend to be sessile, as expected for cells with high c-di-GMP. In B, control cells are more motile than the illuminated cells in A. (C,D) Family trees of a single corresponding family in (A) and (B), where the color corresponds to the same family. Illuminated cells (C) tend to stay adhered across multiple generations, whereas control cells (D) display more surface motility and detachments. See Movie 1 for a representative video of the optogenetic reporter experiment. Figure 5 – Figure supplement 1 shows a schematic of the ATIM apparatus. Figure 5 – See Figure supplement 2 for the data from Figure 5 overlayed onto Figure 4C, showing that ptogenetic-controlled families follow the trend of family behavior observed in wt PAO1 cells.

## Discussion

Collectively, our data show that heterogeneity in cellular levels of c-di-GMP, generated by the Wsp system in response to surface sensing, leads to two distinct physiological subpopulations. Phenotypic heterogeneity of single cells is a common phenomenon in bacteria that is thought to be beneficial at the population level by allowing a single genotype to survive sudden environmental changes and by promoting a division of labor between costly behaviors that support the growth and survival of the population (*26*). Sources of phenotypic heterogeneity include bistability (*27*) and stochasticity (*28*) of gene expression, unequal partitioning of proteins during cell division due to low abundance (*28*), epigenetic modifications resulting in phase variation (*29*), or through asymmetrical cell division (*30*, *31*). In this study, we show that the Wsp system generates heterogeneity in c-di-GMP signaling, and it is never fully activated in 100% of wild-type, surface-attached cells. Moreover, we show that such heterogeneity results in phenotypic changes for entire family lineages of descendent cells. It is interesting that correlations between c-di-GMP, surface motility, and surface detachment probability are strong when considered for an entire lineage in a bacterial family tree, but weak when considered at the individual cell level. This form of correlation suggests that the enforcement of surface sensing outcomes (ex: the activation of DGCs, attenuation of motility) is slow compared to the cells’ division times, and that c-di-GMP signaling is propagated across multiple generations. Additionally, proteins such as DGCs activated by surface sensing may not be passed down to daughter cells equally after division, especially if their number is not large or if they are assymetrically partitioned, which may be one mechanism that leads to the heterogeneity in c-di-GMP levels.

If we overwhelm WspR-generated c-di-GMP heterogeneity by using optogentically-induced sustained c-di-GMP production, we find that phenotypic heterogeneity is lost, and that illuminated cells deterministically become sessile and form microcolonies. Interestingly, our optogenetic experiments show that sustained c-di-GMP production for more than one generation is required before commitment to the sessile lifestyle. This observation is consistent with the fact that we see strong correlations between c-di-GMP levels and motility behavior at the lineage level and not at the individual cell level. Moreover, since the WspR surface sensing system generates heterogeneous c-di-GMP levels, this requirement of sustained c-di-GMP production for more than one generation is inherently difficult for wild-type cells to meet, and virtually guarantees the simultaneous existence of motile and sessile subpopulations. This phenotypic heterogeneity, which has been ‘hardwired’ into the structure of c-di-GMP surface sensing networks, allows for a division of the labor during early biofilm formation, with one subpopulation committing to initiating the protective biofilm lifestyle, while the other subpopulation is free to explore the surface and potentially colonize distant, perhaps more favorable, locations.

## Materials and Methods

### Bacterial strains and growth conditions

The strains, plasmids, and primers used in this study are listed in Table 1. *Escherichia coli* and *P. aeruginosa* strains were routinely grown in Luria–Bertani (LB) medium and on LB agar at 37°C. For the flow cell experiments, *P. aeruginosa* was grown in either LB or FAB minimal medium supplemented with 10mM or 0.6mM glutamate at room temperature (*7*). For flow cytometry experiments, *P. aeruginosa* was grown in either LB medium or in Jensen’s defined medium with glucose as the carbon source (*21*). For the tube biofilm and c-di-GMP measurements, *P. aeruginosa* strains were grown in Vogel-Bonner Minimal Medium (VBMM; (*32*)). Antibiotics were supplied where necessary at the following concentrations: for *E. coli*, 100 µg/mL ampicillin, 10 µg/mL gentamicin, and 10 or 60 µg/mL tetracycline; for *P. aeruginosa*, 300 µg/mL carbenicillin, 100 µg/mL gentamicin, and 100 µg/mL tetracycline. P_*cdrA*_::*gfp*_ASV_ reporter and vector control plasmids were selected with 100 µg/mL gentamicin for *P. aeruginosa* strains and 10 µg/mL gentamicin for *E. coli*.

**Table 1.**
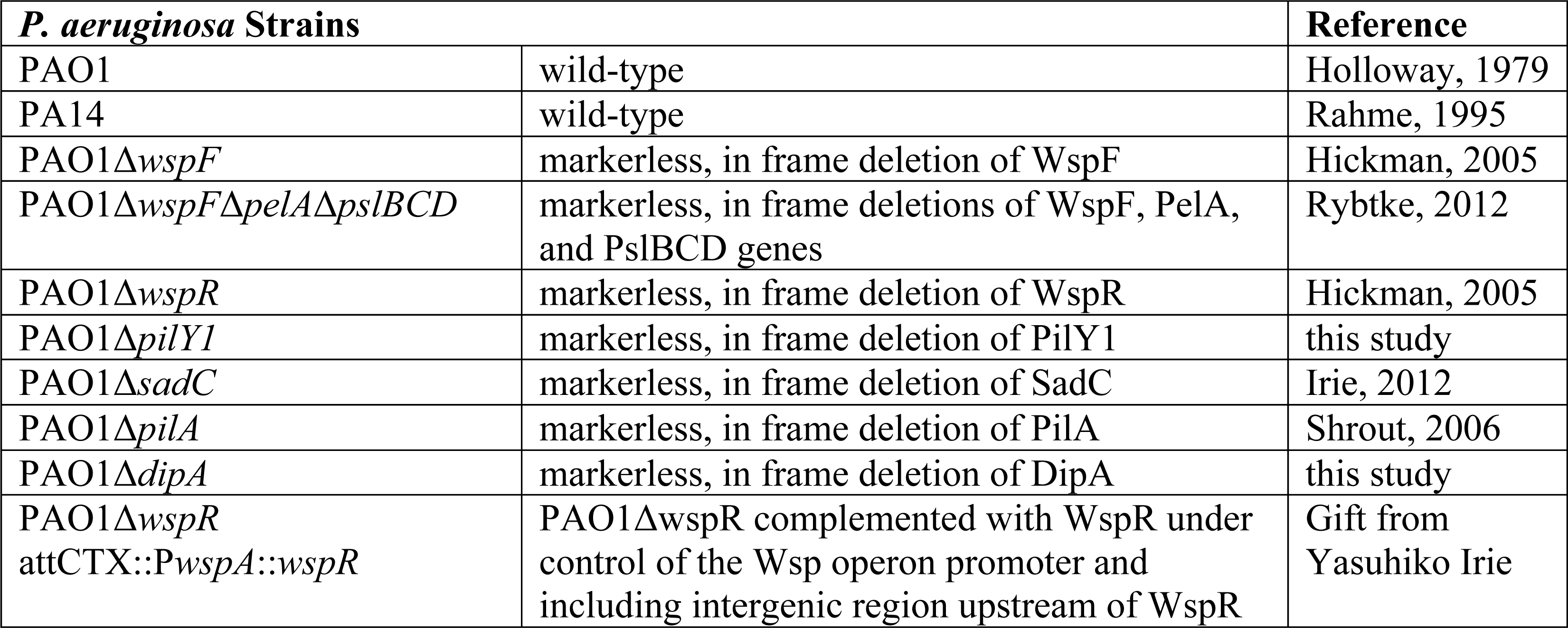
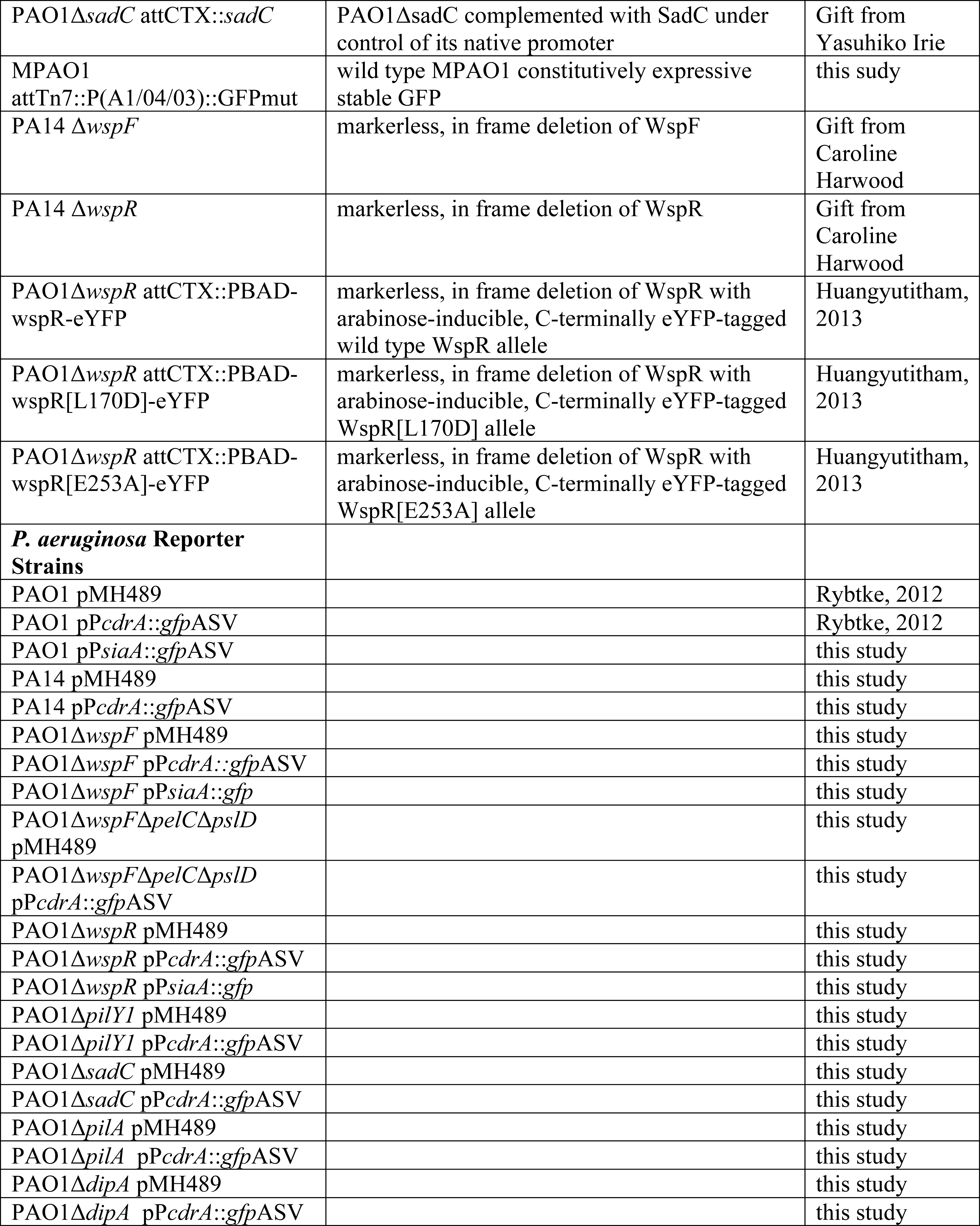
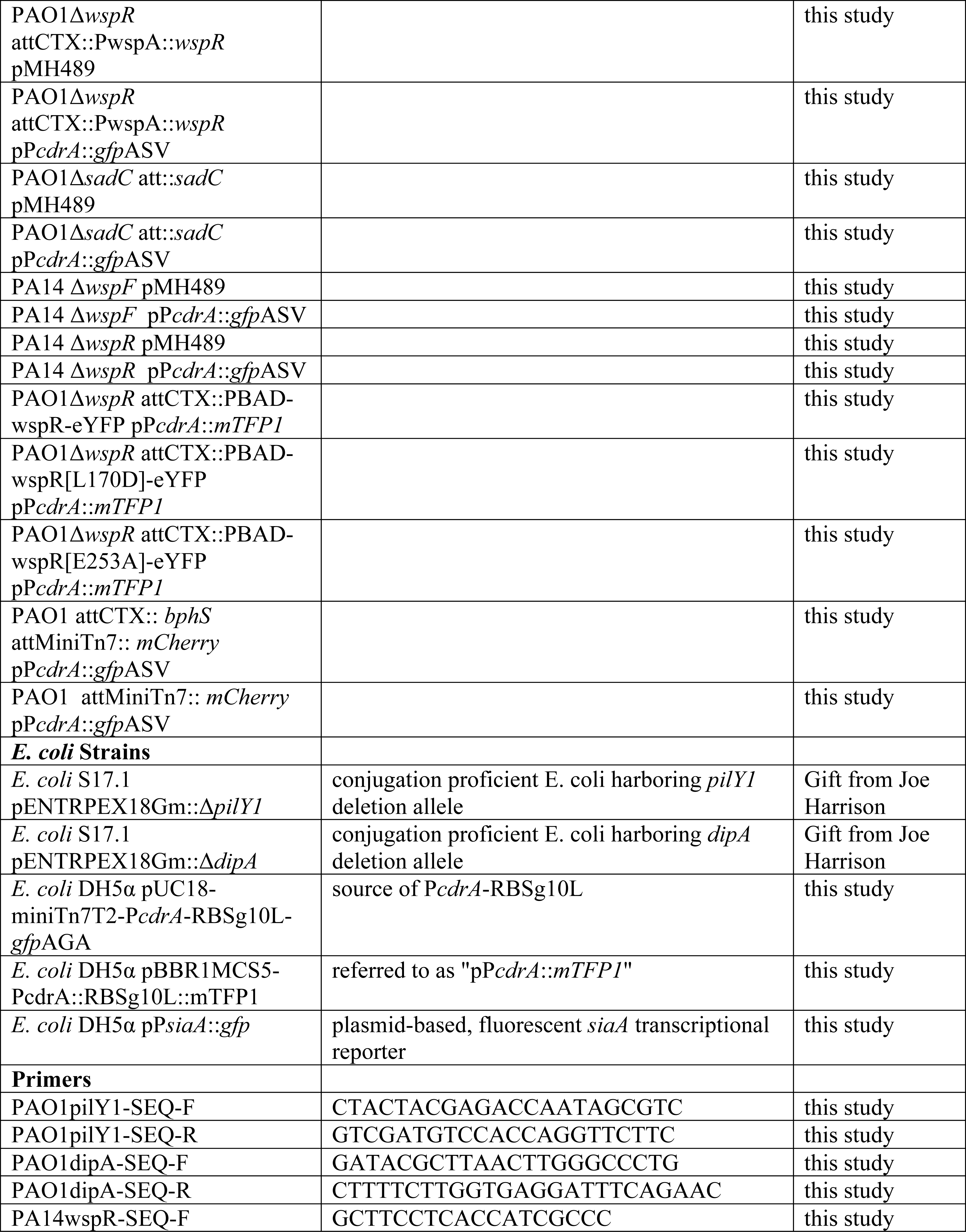
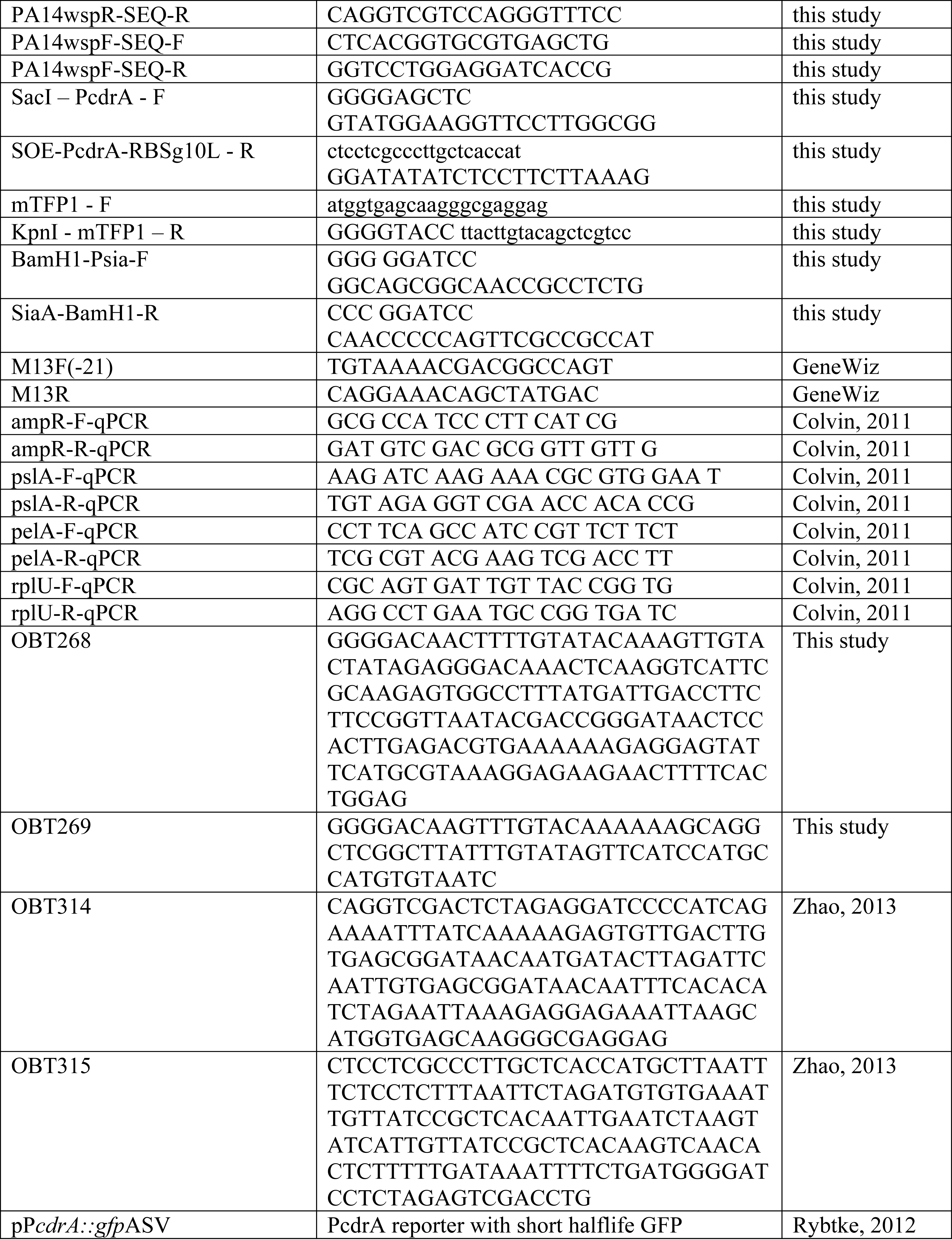
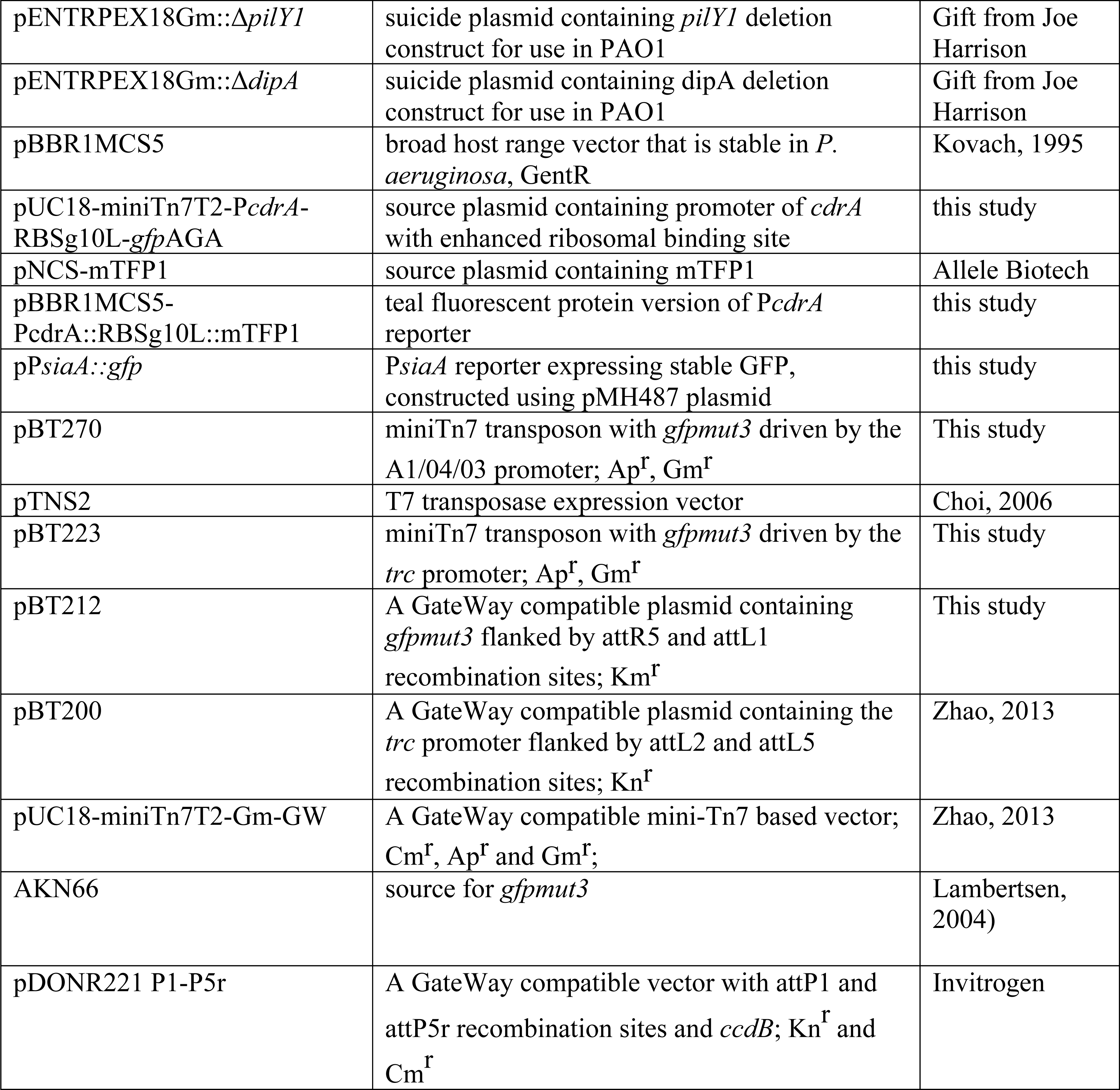
Strains, primers, and plasmids used in this study.

PAO1 Δ*pilY1* was constructed using two-step allelic exchange following conjugation of wild type PAO1 with *E. coli* S17.1 harboring pENTRPEX18Gm::Δ*pilY1* (a gift from Joe Harrison) as previously described (*33*). PAO1 Δ*pilY1* was identified by colony PCR using primers PAO1pilY1-SEQ-F and PAO1pilY1-SEQ-R. PAO1 Δ*dipA* was constructed similarly by conjugation of wild type PAO1 with *E. coli* S17.1 harboring pENTRPEX18Gm::Δ*dipA* (a gift from Joe Harrison). PAO1 Δ*dipA* was identified by colony PCR using primers PAO1dipA-SEQ-F and PAO1dipA-SEQ-R. PA14 Δ*wspR* and Δ*wspF* deletion mutants were confirmed by PCR using primers PA14wspR-SEQ-F and PA14wspR-SEQ-R or PA14wspF-SEQ-F and PA14wspF-SEQ-R, respectively.

To create MPAO1 attTn7::P(A1/04/03)::GFPmut, the miniTn7 from pBT270 was integrated into the chromosome of *P. aeruginosa* PAO1 with the helper plasmid pTNS2, as previously described (*34*). pBT270 was created by introducing the constitutive A1/04/03 promoter (*35*) and removing the trc promoter from pBT223 using the QuikChange Lightning Kit (Agilent Technologies) and the oligonucleotides OBT314 and OBT315. pBT223 was constructed via recombineering of pBT200, pUC18-miniTn7T2-Gm-GW, and pBT212 using Multisite Gateway technology (Invitrogen). pBT212 was constructed by cloning the *gfpmut3* from AKN66 using OBT268 and OBT269, and recombining the PCR product with pDONR221 P1-P5r.

### Construction of optogenetic, c-di-GMP reporter strain in *P. aeruginosa*

Chromosomal insertion of *bphS* was achieved using the mini-CTX system and these strains were marked with different fluorescent proteins by mini-Tn7 site-specific transposition essentially as previously described (*34*, *36*). First, a *bphS* fragment obtained from the plasmid pIND4 was cloned into the vector mini-CTX2 with the *PA1/O4/O3* promoter upstream of the MCS via a two-piece ligation. The constructed plasmid was electroporated into PAO1 and the corresponding recombinant strain was identified by screening on LB agar plates containing 1mM IPTG and 100 µg/mL tetracycline. Then, the strains were electroporated with a pFLP2 plasmid and distinguished on LB agar plates containing 5% (w/v) sucrose for the excision of the resistance marker. The c-di-GMP reporter plasmid and mCherry/EGFP marked *bphS* mutants were constructed as described above. The c-di-GMP reporter plasmid (P_*cdrA*_::*gfp*_ASV_) was electroporated into the mCherry marked *bphS* mutant to monitor the intracellular c-di-GMP level.

### Cyclic di-GMP measurement and qRT-PCR of tube biofilms

Measurement of c-di-GMP in tube biofilm cells was performed as previously described (*4*). Transcriptional analysis of PelA expression in tube biofilms was performed as described in the “FACS and qRT-PCR of c-di-GMP reporter cells” section.

### Crystal violet attachment assays

Crystal violet assays were performed essentially as previously described to measure biofilm biomass, except using gentle washing after 2-6 hours of static incubation (*8*). To measure biofilm biomass at 24 hours, the crystal violet assay was performed as previously described without gentle washing (*37*).

### Flow cell time course experiments and confocal microscopy

*P. aeruginosa* cells harboring the pP_*cdrA*_::*gfp*_ASV_ reporter plasmid or a promotorless vector control (pMH489) were grown to mid-log in LB with 100 µg/mL gentamicin (Gm100) from LB Gm100 plates or from FAB + 10mM glutamate overnight broth cultures in FAB + 10mM glutamate. Mid-log cells were back diluted into 1% LB or FAB + 0.6mM glutamate and flow chambers were inoculated at a final OD_600_ 0.1 and inverted for 10 minutes to allow cells to attach before induction of flow. Clean media was used to wash non-attached cells by flow at 40mL per hour for 20 minutes. Flow was then reduced to a final constant flow rate of 3mL per hour and bacteria were imaged immediately on a Zeiss LSM 510 scanning confocal laser microscope (t=0h). Flow cells were incubated at a constant flow rate at room temperature and imaged hourly for up to 24 hours. For every strain and time point, 5 fields of view and a minimum of 300 cells were captured using identical microscope settings to image GFP fluorescence across all experiments. Images were analyzed using using Volocity software (Improvision, Coventry, UK). Cells were counted as pP_*cdrA*_::*gfp*_ASV_ reporter “on” if their mean GFP fluorescence intensity per pixel was greater than two-fold above the background GFP fluorescence intensity (approximately 340). Data are presented in terms of the percentage of cells with an average GFP fluorescence per pixel twofold more intense compared to the background (pP_*cdrA*_::*gfp*_ASV_ reporter “on”). Microscopy images were artificially colored to display GFP fluorescence as green.

### Construction of pP*siaA*::*gfp*

A region 259 bp upstream through 21 bp into the coding sequence of *siaA* was amplified from PAO1 genomic DNA using primers BamH1-Psia-F and SiaA-BamH1-R, then gel purified using a QIAquick gel extraction kit (Qiagen, Hilden, Germany) digested with BamH1, then column purified with a QIAquick PCR purification kit (Qiagen, Hilden, Germany) to remove BamH1. The GFP expression vector pMH487, which contains the *gfp*mut3 gene with an RNase III splice site and lacking a promoter (*38*), was digested with BamH1, treated with Antarctic phosphatase (New England Biolabs, Ipswich, MA), then column purified with a QIAquick PCR purification kit (Qiagen, Hilden, Germany) to remove BamH1. The P*siaA* allele was ligated into digested pMH487, then transformed into *E. coli* DH5α, purified, and sequenced using primer M13F(−21) (Genewiz). The reporter pP_*siaA*_::*gfp* was electroporated into *P. aeruginosa* as previously described and maintained under gentamycin selection at 100 µg/mL.

### Multi-generation single cell tracking of type IV motility and c-di-GMP reporter activity

Wild type PAO1 harboring the pP_*cdrA*_::*gfp*_ASV_ reporter was grown shaking for 20 hours in FAB media with 6mM glutamate. The flow cell inoculum was prepared by diluting the culture to a final OD_600_ of 0.01 in FAB with 0.6mM glutamate. The flow cell inoculum was injected into the flow cell (Department of Systems Biology, Technical University of Denmark) and allowed to incubate for 10 minutes at 30°C prior to flushing with media at 30mL/h for 10 minutes. Experiments were performed under a flow rate of 3mL/hour for a total of 40 hours.

Images were acquired with an Olympus IX81 microscope equipped with a Zero Drift Correction autofocus system, a 100× oil objective with a 2× multipler lens, and an Andor iXon EMCCD camera using Andor IQ software. Bright-field images were recorded every 3 seconds and GFP fluorescence every 15 minutes. Acquisition continued for a total recording time of 40 hours, which resulted in approximately 48000 bright-field images, and 160 fluorescence images.

Images were analyzed in MATLAB to track bacterial family trees, GFP fluorescence, and surface motility essentially as previously described(*23*) with the following modifications. Image analysis, family tracking and manual validation, family tree plotting, and tree asymmetry λ calculations were performed as previously described(*23*) without modification. GFP fluorescence intensities were normalized by calculating the distribution of intensities per cell per frame (extracted by using the binary image as a mask) and then setting the minimum and maximum intensities to the 1^st^ and 99^th^ percentiles of this distribution for each dataset. I_c-di-GMP_ (relative normalized c-di-GMP reporter intensity) was calculated by averaging the normalized fluorescence intensities across all members of a family. F_motile_ (fraction of time that cells in a family are motile) was calculated as follows. For each family, every cell trajectory in the family was divided into time intervals. For each time interval, presence or absence of motility was determined using a combination of metrics, including Mean Squared Displacement (MSD) slope, radius of gyration, and visit map. MSD slope quantifies the directionality of movement relative to diffusion. Radius of gyration and visit map are different metrics for quantifying the average distance traveled on the surface. F_motile_ was then calculated by the fraction of these time intervals that have motility. This calculation was modified from the “TFP activity metric” previously described(*23*).

### Setup of Adaptive Tracking Illumination Microscopy

Figure 5 – Supplement 1 shows a schematic of the Adaptive Tracking Illumination Microscopy (ATIM) setup. An inverted fluorescent microscope (Olympus, IX71) was modified to build the ATIM. The modification includes: 1) a commercial DMD-based LED projector (Gimi Z3) was used to replace the original bright-field light source, in which the original lenses in the projector were removed and three-colored (RGB) LEDs were rewired to connect to an external LED driver (ThorLabs) controlled by a single chip microcomputer (Arduino UNO r3); 2) the original bright-field condenser was replaced with an air objective (40× NA = 0.6, Leica); and 3) an additional 850 nm LED light (ThorLabs) was coupled to the illumination optical path using a dichroic mirror (Semrock) for the bright-field illumination. Note that 850 nm LED light is safe light to ensure that the bright-filed illumination does not affect optogenetic manipulation. The inverted fluorescent microscope (Olympus, IX71) equipped with a 100×oil objective and a sCMOS camera (Zyla 4.2 Andor) was used to collect bright-field images with 0.2 frame rate. The bright-field images were further analyzed to track multiple single cells in real time using a high-throughput bacterial tracking algorithm coded by Matlab. The projected contours of selected single cells were sent to the DMD (1280 × 760 pixels) that directly controlled by a commercial desktop through a VGA port. The manipulation lights were generated by the red-color LED (640 nm), and were projected on the single selected cells in real time through the DMD, a multi-band pass filter (446/532/646, Semrock) and the air objective. Our results indicated that feedback illuminations could generate projected patterns to exactly follow the cell movement (Figure 5 – Supplemental 1B) or single cells divisions (Figure 5 – Supplemental 1C) in real time.

### Manipulation of c-di-GMP expression in single initial-attached cells

The bacterial strain PAO1-*bphS*-P*cdrA*-GFP-mCherry was inoculated into a flow cell (Denmark Technical University) and continuously cultured at 30.0 ± 0.1°C by flowing FAB medium (3.0 mL/h). The flow cell was modified by punching a hole with a 5 mm diameter into the channel, and the hole was sealed by a coverslip that allows the manipulation light to pass through. An inverted fluorescent microscope (Olympus, IX71) equipped with a 100× oil objective and a sCMOS camera (Zyla 4.2 Andor) was used to collect bright field or fluorescent images with 0.2 or 1/1800 frame rate respectively. The power density of the manipulation lights was determined by measuring the power at the outlet of the air objective using a power meter (Newport 842-PE). GFP or mCherry was excited using a 480 nm or 565 nm LED lights (ThorLabs) and imaged using single-band emission filters (Semrock): GFP (520/28 nm) or mCherry (631/36 nm). Initial-attached cells were selected to be manipulated using ATIM with the illumination at 0.05 mW/cm^2^, which allowed us to compare the results arising from illuminated or un-illuminated mobile cells in one experiment. The c-di-GMP levels in single cells were gauged using the ratio of GFP and mCherry intensities.

### Lectin staining and flow cytometry

Glass culture tubes were inoculated with 1mL of *P. aeruginosa* in LB or Jensen’s minimal media at an OD_600_ 0.8 and incubated statically at 37°C for 4 hours. Non-adhered cells were removed by washing three times with 2mL sterile phosphate buffered saline (PBS). Biofilm cells were harvested by vortexing in 1mL PBS with fluorescein-labeled lectins (WFL lectin (100 µg/mL; Vector Laboratories) for Pel, TRITC-labeled HHA (100 µg/mL; EY Laboratories) for Psl) and incubated on ice for 5 minutes. Cells were washed 3 times to remove non-adhered lectin, resuspended in PBS, and immediately analyzed for GFP and TRITC fluorescence on a BD LSRII flow cytometer (BD Biosciences). Events were gated based on forward and side scatter to remove particles smaller than a single *P. aeruginosa* cell and large aggregates.

We used PAO1 cells that did not express GFP (wild type PAO1; Figure 1 – Supplement 3A) or constitutively expressed GFP (PAO1 Tn7::P(A1/04/03)::GFPmut; Figure 1 – Supplement 3B) to define a gate for high GFP fluorescence. We validated this gate using a strain in which we expect very high levels of reporter activity (surface grown PAO1 Δ*wspF*Δ*pelA*Δ*pslBCD* harboring pP*cdrA*::*gfp*_ASV_) and saw that 91.6% of cells had high GFP levels (Figure 1 – Supplement 3C), in agreement with our flow cell characterization of this strain (Figure 2A). We determined gating for TRITC using cells that had not been stained with TRITC-conjugated lectin (Figure 1 – Supplement 5A), as well as two strains that overproduced either Psl (Figure 1 – Supplement 5B) or Pel (Figure 1 – Supplement 5C) that were stained with the appropriate TRITC-conjugated lectin. Our flow cytometry gating procedure accurately gated 99.7% of wild type PAO1 cells (without the P*cdrA* reporter or lectin-staining) as low GFP and low TRITC (Figure 1 – Supplement 5D).

### FACS and qRT-PCR of c-di-GMP reporter cells

Static biofilm reporter cells were grown as described above and harvested without lectin staining. Cells were fixed with 6% paraformaldehyde for 20 minutes on ice, then rinsed once with sterile PBS prior to analysis with a FACSAriaII (BD Biosciences, San Jose, CA). Events were gated first to remove debris and large cellular aggregates, and then gated into cells with low and high GFP fluorescence intensity. The low GFP gate was drawn using wild type PAO1 cells without the gfp gene (Figure 1 – Supplement 6A) and the high GFP gate was drawn using both PAO1 Tn7::P(A1/04/03)::GFPmut (Figure 1 – Supplement 6B) and PAO1 Δ*wspF* Δ*pelA* Δ*pslBCD* P_*cdrA*_::*gfp*_ASV_ reporter (Figure 1 – Supplement 6C). As expected, wild type PAO1 pP_*cdrA*_::*gfp*_ASV_ reporter cells that had been harvested after 4 hours of surface attachment to glass in static LB liquid culture displayed subpopulations of high GFP, reporter “on” cells (30.8% of the population) and “off” (57.2%) cells (Figure 1 – Supplement 6D), whereas this same strain grown to mid-log planktonically in LB displayed mostly reporter “off” cells (Figure 1 – Supplement 6E). Cells were sorted at 4°C by flow assisted cell sorting (FACS) to collect 100,000 events into TRIzol LS (Thermo Fisher Scientific, Waltham, MA). RNA was extracted from sorted cells by boiling immediately for 10 minutes and following the manufacturer’s instructions for RNA isolation. DNA was digested by treating with RQ1 Dnase I (Promega, Madison, WI) and samples were checked for genomic DNA contamination by PCR to detect *rplU*. Expression of *pelA*, *pslA*, and *ampR* was measured by quantitative Reverse Transcriptase PCR (qRT-PCR) using the iTaq Universal SYBR Green One-Step kit (Biorad, Hercules, CA) and a CFX96 Touch Real-Time PCR detection system (Bio- Rad, Hercules, CA). The ΔΔC_q_ was calculated for 3 independent samples of sorted wild type PAO1 P_*cdrA*_::*gfp*_ASV_ reporter biofilm cell populations by normalizing PelA and PslA to relative levels of AmpR expression. Data were presented as the average fold change in PelA or PslA expression in the P_*cdrA*_::*gfp*_ASV_ sorted “on” population (high GFP) relative to the “off” population (low GFP) for the three biological replicates.

### WspR-YFP foci and pP*cdrA*::*mTFP1* reporter

A version of the pP*cdrA* reporter was constructed in the pBBR1MCS5 plasmid to express mTFP1 instead of GFP, for use with YFP-tagged WspR proteins. The P*cdrA* promoter and an enhanced ribosomal binding site from the gene 10 leader sequence of the T7 phage (g10L) was amplified from pUC18-miniTn7T2-P_*cdrA*_-RBSg10L-*gfp*_AGA_ using primers SacI-PcdrA-F and SOE-PcdrA-RBSg10L-R. The primers mTFP1-F and KpnI-mTFP1-R were used to amplify the mTFP1 gene from plasmid pNCS-mTFP1 (Allele Biotech, San Diego, CA). The P_*cdrA*_::RBSg10L::*mTFP1* allele was constructed by SOE-PCR using primers SacI-PcdrA-F and Kpn1-mTFP1-R, then pBBR1MCS5 and the SOE PCR product were doubly digested with SacI/KpnI. Digested pBBR1MCS5 was treated with Antarctic phosphatase, then both digests were gel purified and ligated. The ligation was transformed into *E. coli* DH5α, and plasmid from clones growing on LB with 10 µg/mL gentamycin were sequenced with primers M13F and M13F(−21) (GeneWiz). Fluorescence of the pP_*cdrA*_::*mTFP1* reporter was measured in Wsp mutants in a fluorimeter (BioTek Synergy H1 Hybrid Reader, BioTek Instruments, Inc., Winooski, VT, USA) and in flow cells to confirm its activity resembled that of pP_*cdrA*_::*gfp*_ASV_. The pP*cdrA*::*mTFP1* reporter was electroporated into *P. aeruginosa* strains with the native WspR deleted and harboring an arabinose-inducible copy of WspR-YFP on its chromosome (*12*). Cells were grown on LB agar plates with 100 µg/mL gentamycin and 1% arabinose for 10 hours, then transferred to an agar pad for imaging. WspR-YFP foci and mTFP1 fluorescence was imaged using a Nikon Ti-E inverted wide-field fluorescence microscope with a large-format scientific complementary metal-oxide semiconductor camera (sCMOS; NEO, Andor Technology, Belfast, United Kingdom) and controlled by NIS-Elements. WspR-YFP foci were detected as previously described (*12*).

## Acknowledgements

We thank Drs. Julie Cass and Paul Wiggins providing the wide-field microscope and cMOS camera to image WspR-eYFP clusters, Drs. Joe J. Harrison and Yasuhiko Irie for the gift of bacterial strains, and Dr. Keiji Murakami for performing c-di-GMP measurements.

---

**Figure 1 – Figure supplement 1.**
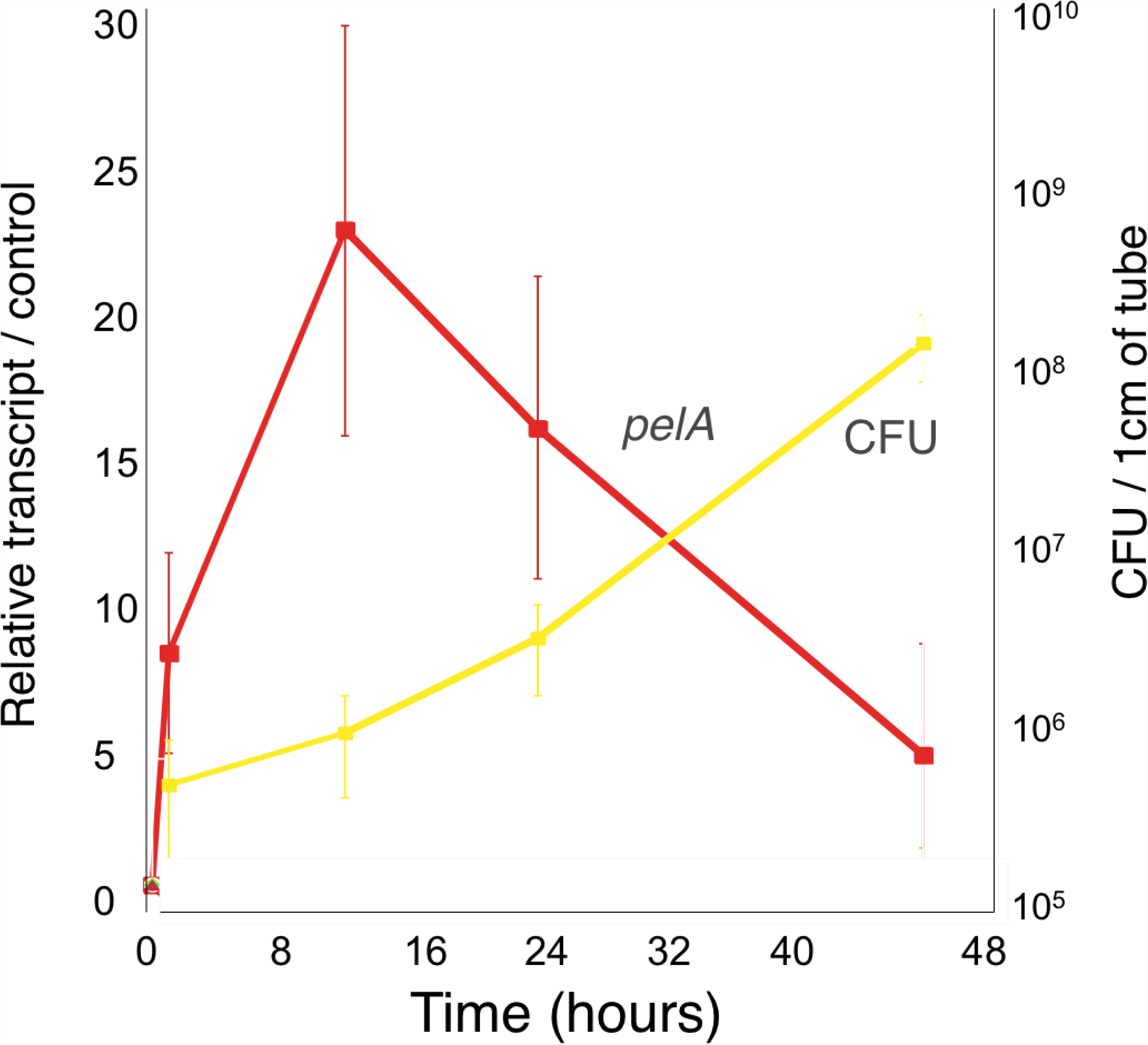
The c-di-GMP-regulated promoter of the Pel polysaccharide operon is transcriptionally activated almost 10-fold compared to planktonic cells within 30 minutes of attachment of PAO1 to a silicone tube. qRT-PCR was performed to detect *pelA* transcript levels in silicone tube biofilm cells compared to planktonic cells (red). Biofilm transcript levels were normalized to planktonic levels at each time point. Colony forming units (CFU) of biofilm attached cells is plotted in yellow at each time point.

**Figure 1 – Figure supplement 2.**
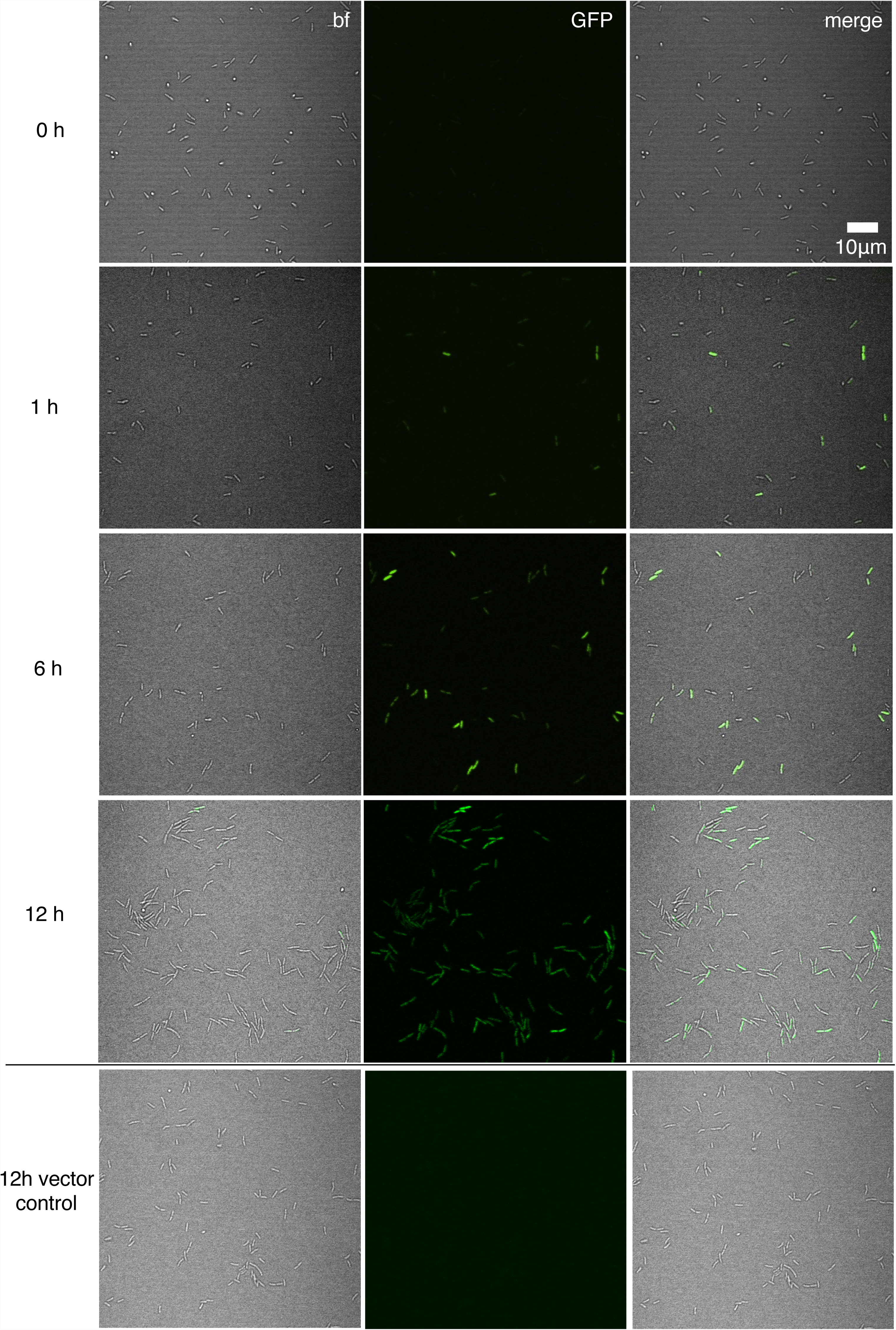
Representative time course images showing the c-di-GMP reporter (P_*cdrA*_::*gfp*_ASV_) transitioning from inactive upon initial attachment of wild type PAO1 (0 hr) to active in a subpopulation of cells between 1 and 12 hours during a flow cell experiment. A strain harboring the vector control, which encoded *gfp*_ASV_, but lacks the P_*cdrA*_ promoter was imaged alongside each reporter strain (in the appropriate genetic background) to confirm that GFP fluorescence was not due to random expression from the plasmid. Wild type PAO1 P_*cdrA*_::*gfp*_ASV_ was grown in 1% LB and imaged by CSLM. bf = bright field, merge = bright field and GFP channels combined.

**Figure 1 – Figure supplement 3.**
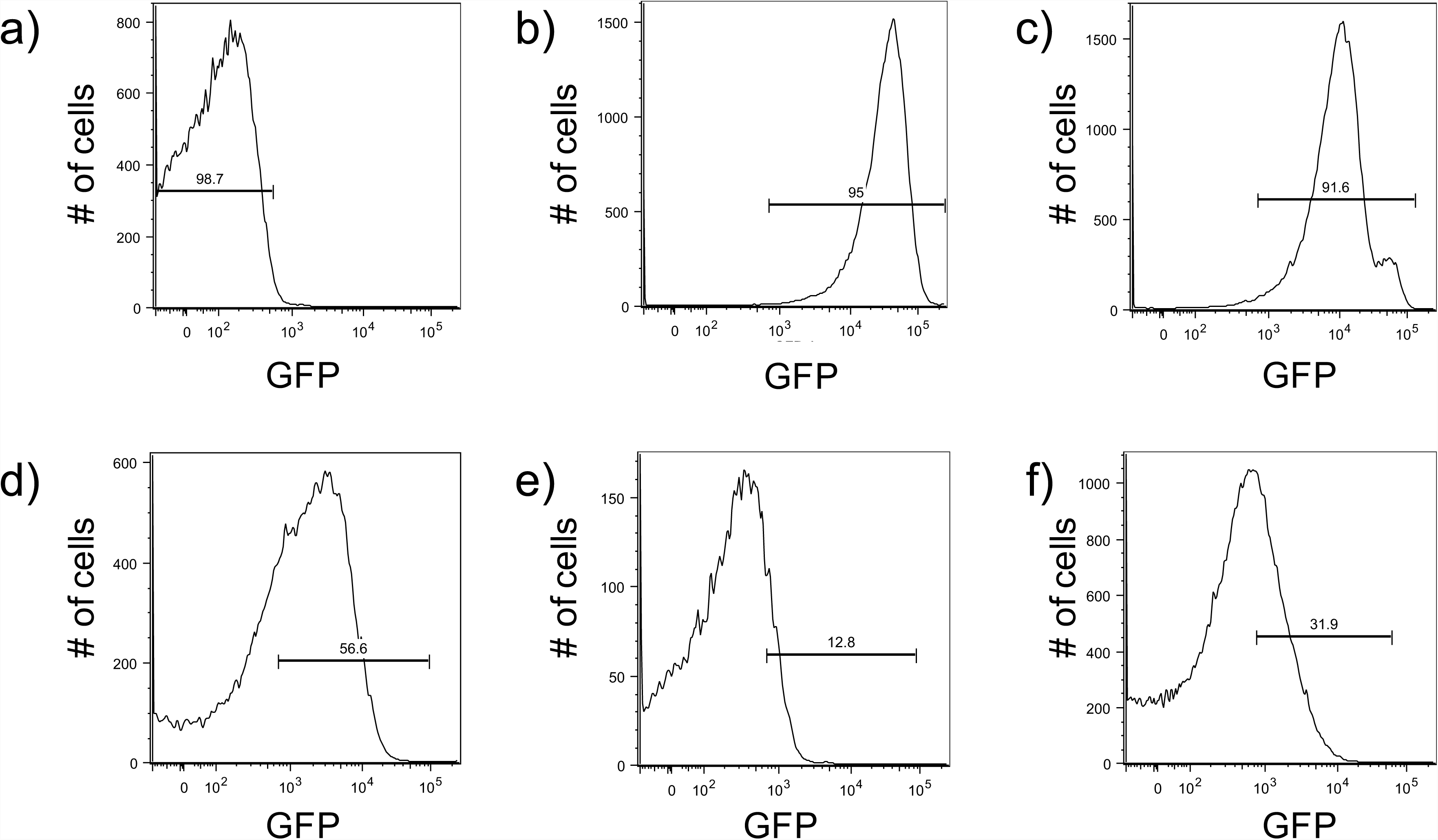
Development of a protocol to monitor pP_*cdrA*_::*gfp*_ASV_ using an LSRII flow cytometer. Brackets indicate gates for “on” (GFP above 1.7 x 10^2^ RFU) or “off” (GFP below 1.7 x 10^2^ RFU) reporter cells and the number above the bracket indicates the percentage of cells that fall within that gate. a) Wild type PAO1 cells were used to determine the background level of fluorescence on the BD Aria III for GFP measurements. The population of cells falls below 10^3^ RFU. b) A *P. aeruginosa* strain constitutively expressing stable GFP (PAO1 Tn7::P(A1/04/03)::GFPmut) was used to determine gating for cells with high GFP, with the population ranging from 10^3^ to 10^5^ RFU. c) Surface grown PAO1 Δ*wspF*Δ*pel*Δ*psl* harboring the pP_*cdrA*_::*gfp*_ASV_ was used to validate the gate for collection of cells with high reporter activity (10^3^ to 10^5^ RFU; 91.6% of the population). d) Example of gating for reporter “on” cells from wild type PAO1 pP_*cdrA*_::*gfp*_ASV_ cells that had been attached to glass in LB medium for 4 hours. Approximately 56.6% of the population falls into the reporter “on” population. e) Example of gating for “on” cells from wild type PA14 pP_*cdrA*_::*gfp*_ASV_ cells that had been attached to glass in LB medium for 4 hours prior to FACS sorting. Approximately 12.8% of the population falls into the reporter “on” population. f) Example of gating for “on” cells from wild type PA14 pP_*cdrA*_::*gfp*_ASV_ cells that had been attached to glass in Jensen’s media (a condition in which Pel is more abundantly produced than in LB) for 4 hours. Approximately 31.9% of the population falls into the reporter “on” population.

**Figure 1 – Figure supplement 4.**
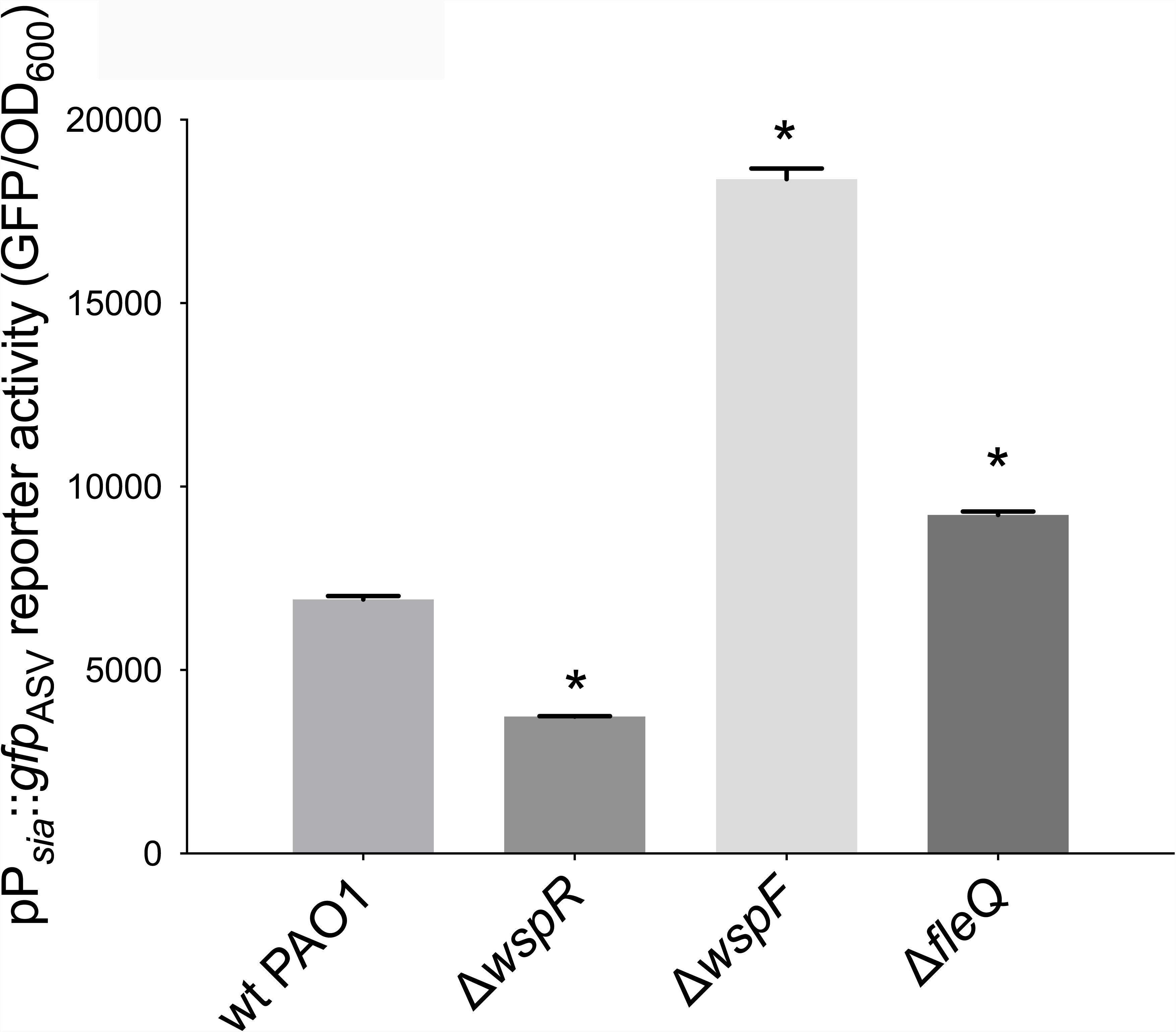
The *siaA* promoter, regulated by elevated c-di-GMP/FleQ is responsive to Wsp-dependent changes in cellular levels of c-di-GMP. Wild type PAO1 and PAO1 mutants harboring the pP_*siaA*_::*gfp* reporter were grown for 20 hours on an LB agar surface with 100 µg/mL gentamycin, resuspended in PBS, and their GFP fluorescence and absorbance at OD_600_ was measured immediately in a spectrophotometer. Asterisk indicates a statistically significant difference in pP_*siaA*_::*gfp* reporter activity relative to wild type PAO1 (p < 0.05, N = 6, ANOVA with post-hoc Dunnett).

**Figure 1 – Figure supplement 5.**
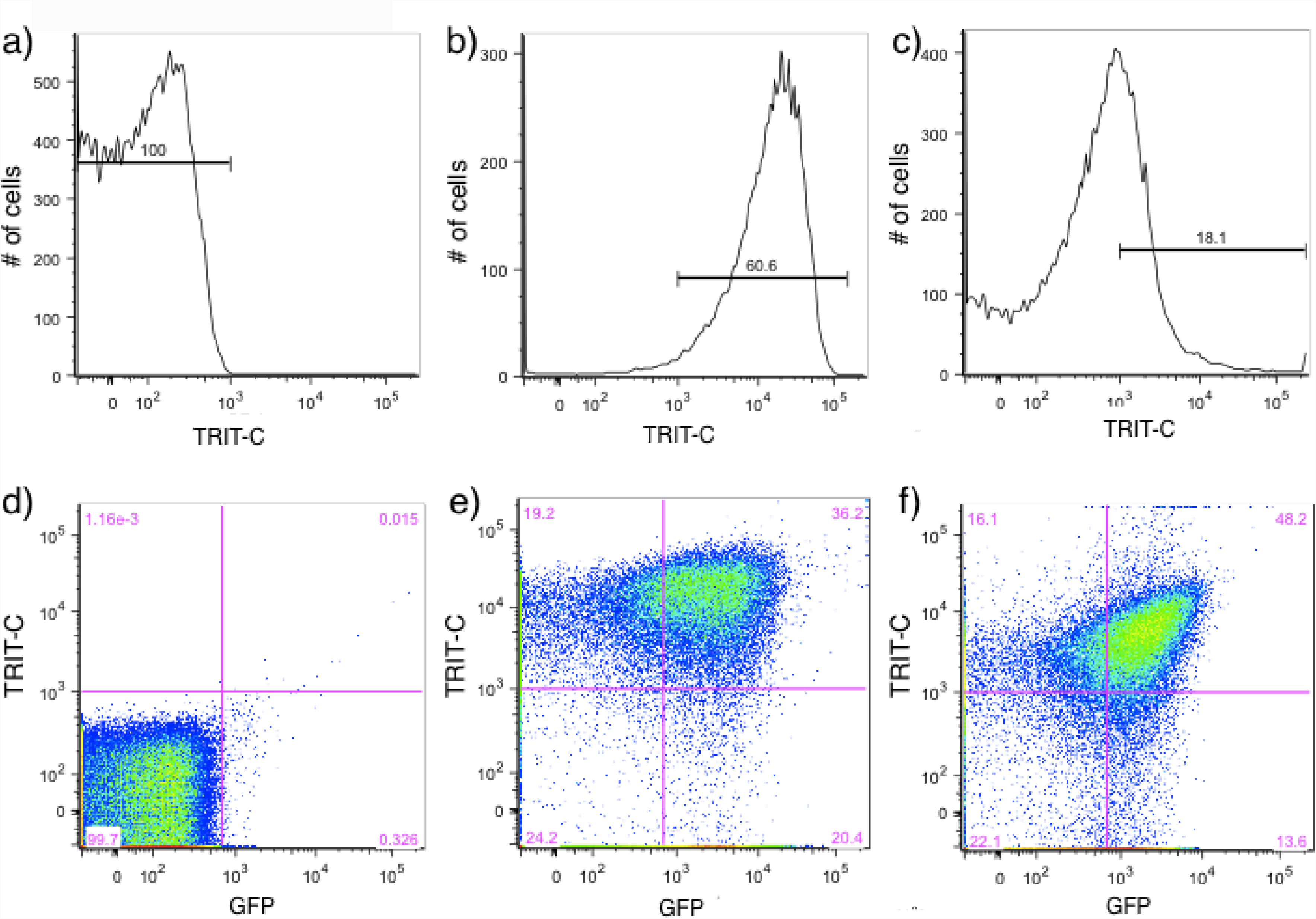
Psl and Pel polysaccharide production is highest in cells with high c-di-GMP as measured by the pP_*cdrA*_::*gfp*_ASV_ reporter. Brackets indicate gates for polysaccharide-producing, TRITC lectin bound cells (TRITC above 10^3^ RFU) or cells without lectin bound (TRITC below 10^3^ RFU) on an LSR II flow cytometer and numbers above the brackets indicate the percentage of total cells that fall within the gate. a) Determination of the background level of TRITC autofluorescence in wild type PAO1 that had not been stained with a TRITC-conjugated lectin. b) Determination of the a high level of TRITC-conjugated Psl-specific lectin binding in PAO1 P_BAD_-psl grown in shaken liquid culture for 4 hours with 1% arabinose before staining with the TRITC-HHA lectin and extensive washing. Approximately 60% of the cells have TRIT-C-HHA lectin bound to their surface. c) Determination of the a high level of TRITC-conjugated Pel-specific lectin binding in PAO1 P_BAD_-*pel* grown in shaken liquid culture for 4 hours with 1% arabinose before staining with the TRITC-WFL lectin and extensive washing. Approximately 18.1% of the cells have TRITC-HHA lectin bound to their surface. d) A scatterplot of pP_*cdrA*_::*gfp*_ASV_ reporter activity (GFP) on the x axis and Psl production (as measured by binding of TRITC-HHA lectin to the cell surface) demonstrating the specificity of GFP and TRITC gating in a negative control condition. Wild type PAO1 cells that do not contain the reporter and had not been stained with lectin mostly fall into the lower left quadrant of “off” GFP and no TRITC-lectin binding. The purple numbers in each quadrant represent the percentage of the total population that falls within that quadrant. Quadrants were drawn based on gating of pP_*cdrA*_::*gfp*_ASV_ reporter activity “on” vs. “off” and lectin bound vs. unbound. e) Psl polysaccharide production is enriched in the population of cells with high c-di-GMP. Representative scatterplot of reporter activity versus Psl lectin binding in wild type PAO1 harboring the pP_*cdrA*_::*gfp*_ASV_ reporter grown for four hours in LB before surface attached cells were harvested, lectin stained, washed, and counted by flow cytometry. f) Pel polysaccharide production is enriched in the population of cells with high c-di-GMP. Representative scatterplot of reporter activity versus Pel lectin binding in wild type PA14 harboring the pP_*cdrA*_::*gfp*_ASV_ reporter grown for four hours in Jensen’s minimal media plus glucose before surface attached cells were harvested, lectin stained, washed, and counted by flow cytometry.

**Figure 1 – Figure supplement 6.**
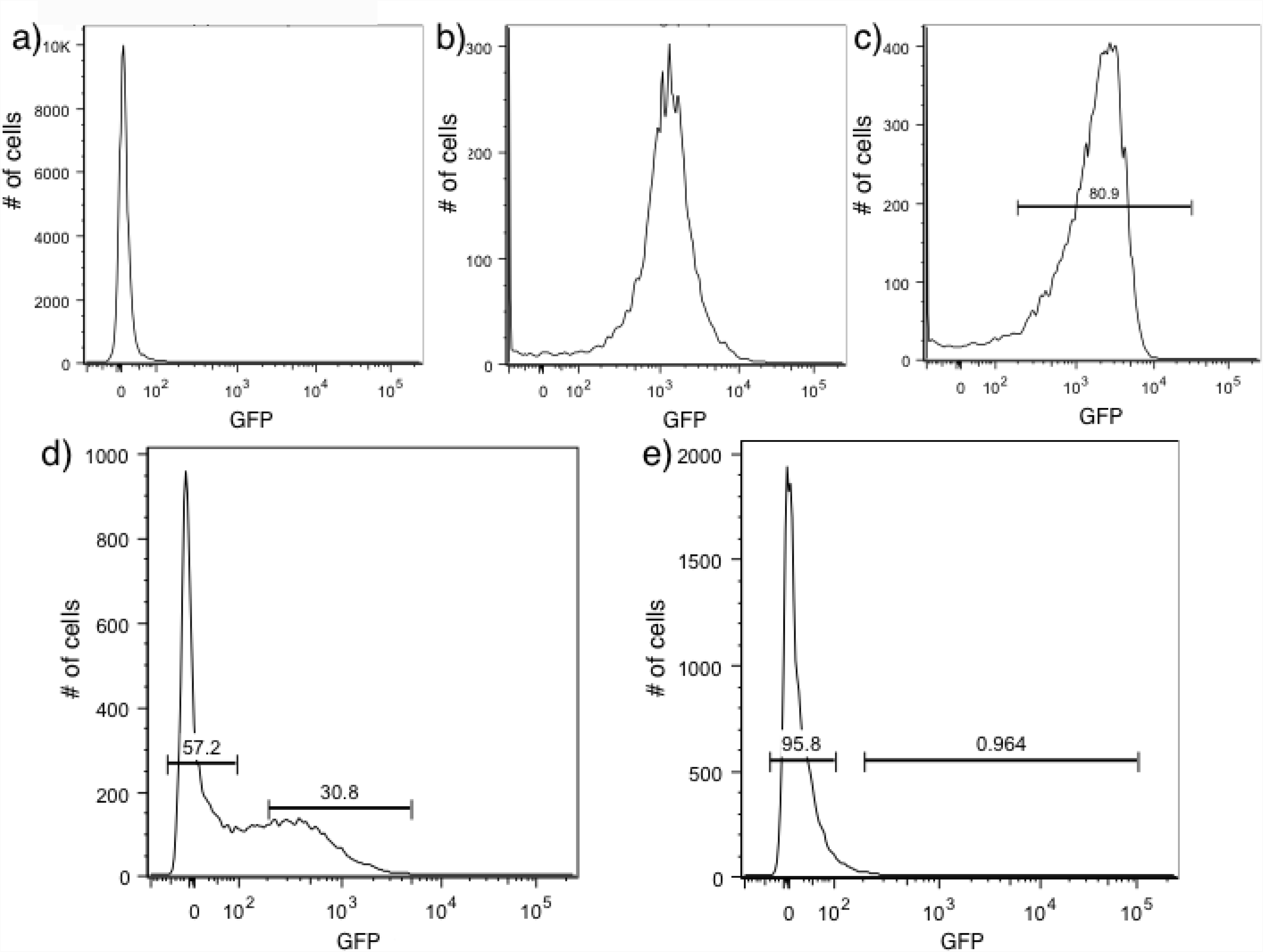
Development of a protocol to sort biofilm cells by pP_*cdrA*_::*gfp*_ASV_ using flow assisted cell sorting (FACS). Brackets indicate gates for “on” (GFP above 1.1 x 10^2^ RFU) or “off” (GFP below 10^2^ RFU) reporter cells on a BD Aria III flow cytometer and the number above the bracket indicates the percentage of cells that fall within that gate. a) Wild type PAO1 cells were used to determine the background level of fluorescence on the BD Aria III for GFP measurements. The population of cells centers around zero RFU. b) A *P. aeruginosa* strain constitutively expressing stable GFP (PAO1 Tn7::P(A1/04/03)::GFPmut) was used to determine gating for cells with high GFP, with the population centering around 10^3^ RFU. c) PAO1 Δ*wspF*Δ*pel*Δ*psl* harboring the pP_*cdrA*_::*gfp*_ASV_ was used to draw a gate for collection of cells with high reporter activity (80% of the population). d) Example of gating for reporter “off” and “on” cells from wild type PAO1 pP_*cdrA*_::*gfp*_ASV_ cells that had been grown on a surface for 4 hours prior to FACS sorting. Approximately 30% of the population falls into the reporter “on” population. A gap was left between the sorted “off” and “on” populations to increase the stringency of the sorting. e) Example of the “off” and “on” gates drawn on wild type PAO1 pP_*cdrA*_::*gfp*_ASV_ cells that were grown planktonically to mid-log, demonstrating that the surface dependent nature of P*cdrA* reporter activity can be detected by flow cytometry.

**Figure 2 – Figure supplement 1.**
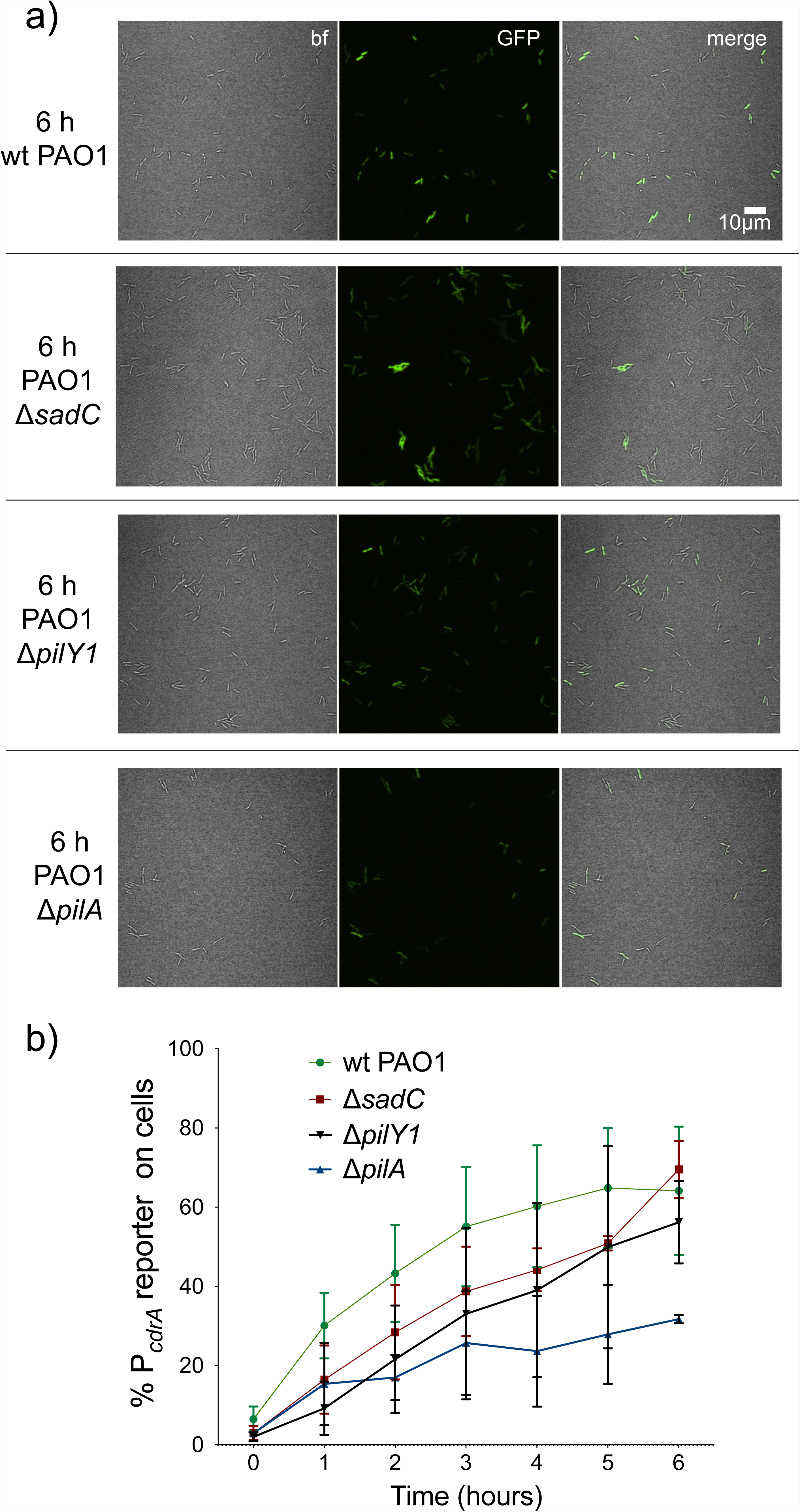
Mutants predicted to inactivate the Pil-Chp surface sensing system largely retain pP_*cdrA*_::*gfp*_ASV_ reporter activity during the first six hours of surface sensing. a) Representative images from wild type PAO1, PAO1 Δ*sadC*, PAO1 Δ*pilY1*, and PAO1 Δ*pilA* after 6 hours of surface attachment. bf = bright field, merge = bright field and GFP channels combined. b) Six hour time course plot of the average percentage of cells from either wild type PAO1 (green), PAO1 Δ*sadC* (red), PAO1 Δ*pilY1* (black), or PAO1 Δ*pilA* (blue) in which the pP_*cdrA*_::*gfp*_ASV_ reporter had turned “on” at each hour. PAO1 Δ*pilA* is significantly different from wild type PAO1 from 2-6 hours (T-test, p < 0.05). Cells were identified as “on” if their average GFP fluorescence was greater than twice the average background GFP fluorescence of the image. Plotted values are the mean of at least 3 biological replicates and error bars are standard deviation.

**Figure 2 – Figure supplement 2.**
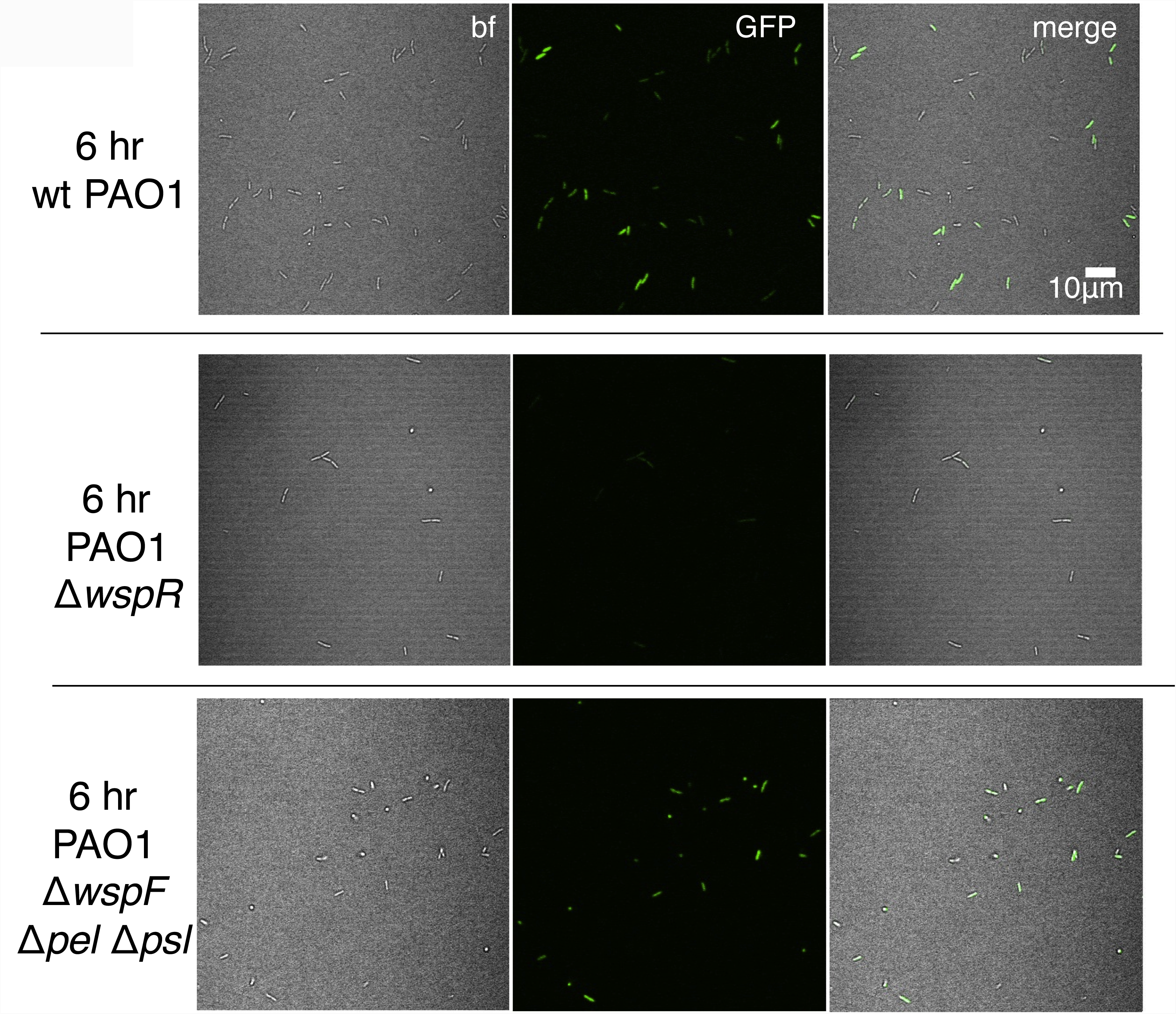
The pP_*cdrA*_::*gfp*_ASV_ reporter is sensitive to Wsp-dependent variation in c-di-GMP during surface sensing. Representative images from wild type PAO1, PAO1 Δ*wspR*, and PAO1 Δ*wspF*Δ*pelC*Δ*pslD* after 6 hours of surface attachment. bf = bright field, merge = bright field and GFP channels combined.

**Figure 2 – Figure supplement 3.**
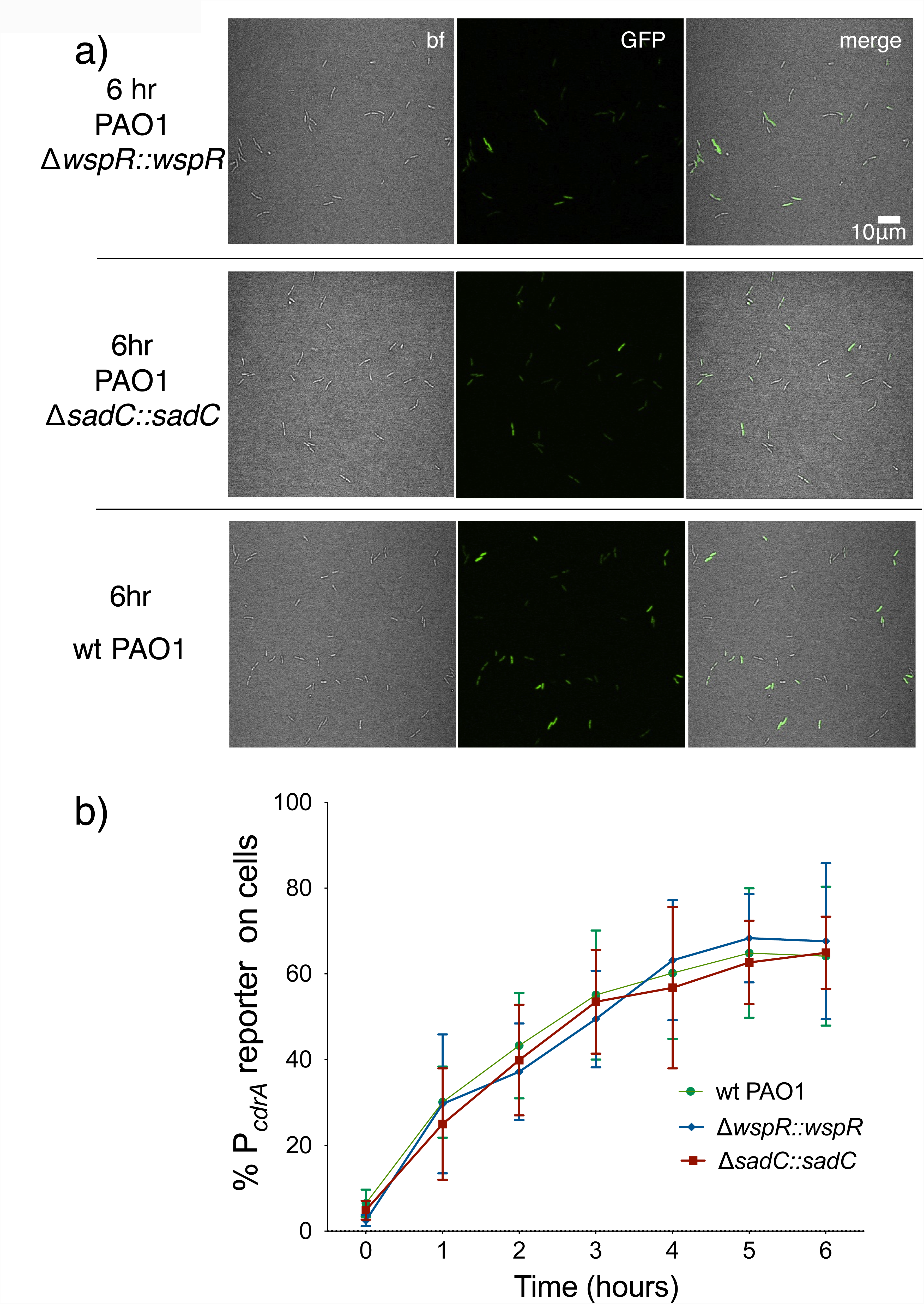
Complemented diguanylate cyclase mutants display wild type levels of P*cdrA*::*gfp*_ASV_ reporter activity. PAO1 Δ*wspR* and PAO1 Δ*sadC* were complemented at a neutral site on the chromosome under control of their native promoters. a) Representative images from wild type PAO1, PAO1 Δ*wspR* attCTX::*wspR* (Δ*wspR::wspR*), and PAO1 Δ*sadC* Tn7::*sadC* (PAO1 Δ*sadC::sadC*) after 6 hours of surface attachment. bf = bright field, merge = bright field and GFP channels combined. b) Six hour time course plot of the average percentage of cells from either wild type PAO1 (green), PAO1 Δ*wspR* attCTX::*wspR* (blue), or PAO1 Δ*sadC* Tn7::*sadC* in which the pP_*cdrA*_::*gfp*_ASV_ reporter had turned “on” at each hour. Cells were identified as “on” if their average GFP fluorescence was greater than twice the average background GFP fluorescence of the image. Error bars = standard deviation, n ≥ 3 biological replicates.

**Figure 2 – Figure supplement 4.**
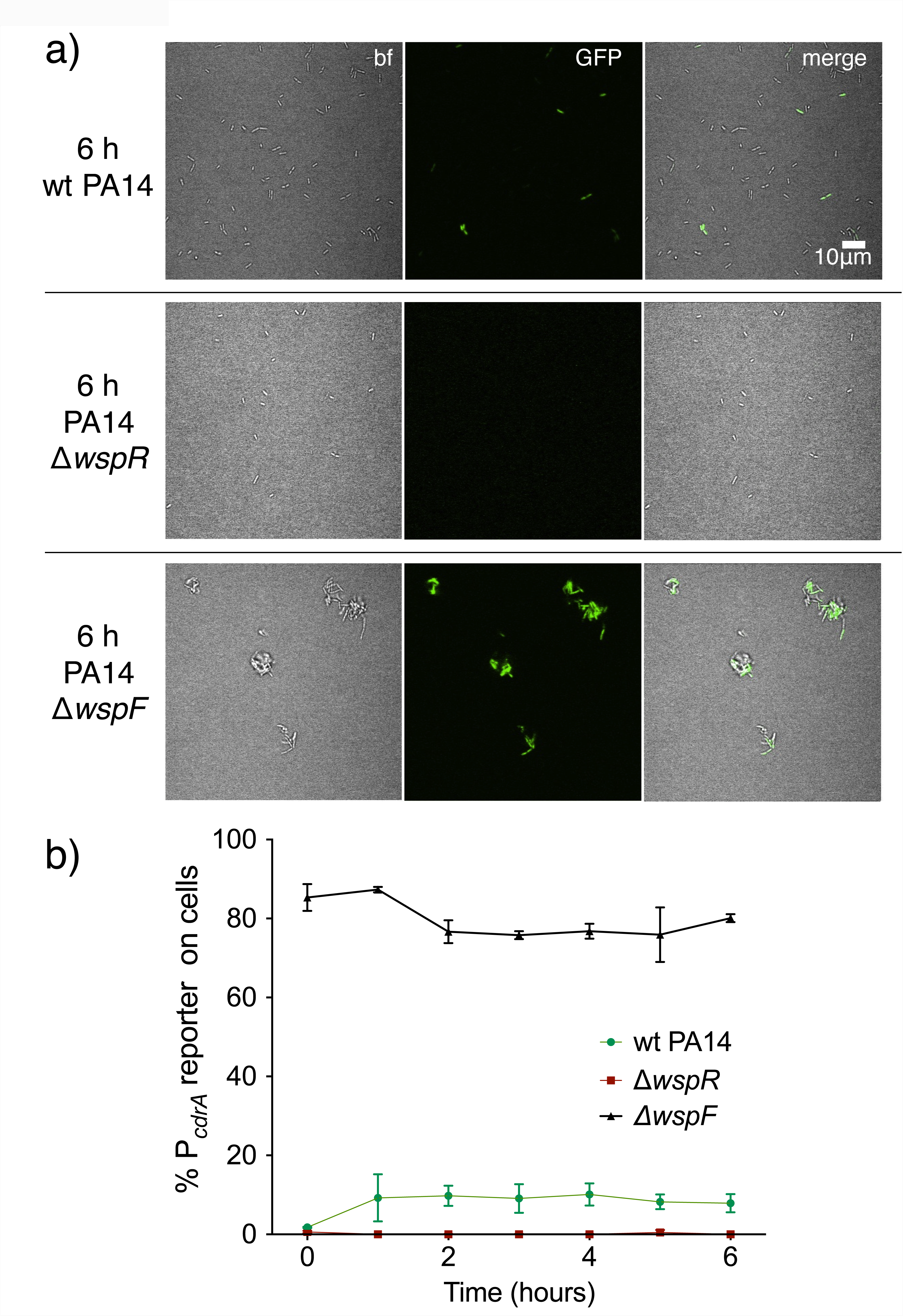
Activity of the pP_*cdrA*_::*gfp*_ASV_ reporter in strain PA14 is dependent on the Wsp system. a) Representative images from wild type PA14, PA14 Δ*wspR*, and PA14 Δ*wspF* after 6 hours of surface attachment. bf = bright field, merge = bright field and GFP channels combined. b) Six hour time course plot of the average percentage of cells from either wild type PA14 (green), PA14 Δ*wspR* (red), or PA14 Δ*wspF* (black) in which the pP_*cdrA*_::*gfp*_ASV_ reporter had turned “on” at each hour. Cells were identified as “on” if their average GFP fluorescence was greater than twice the average background GFP fluorescence of the image. Error bars = standard deviation, n ≥ 3 biological replicates.

**Figure 5 – Figure supplement 1.**
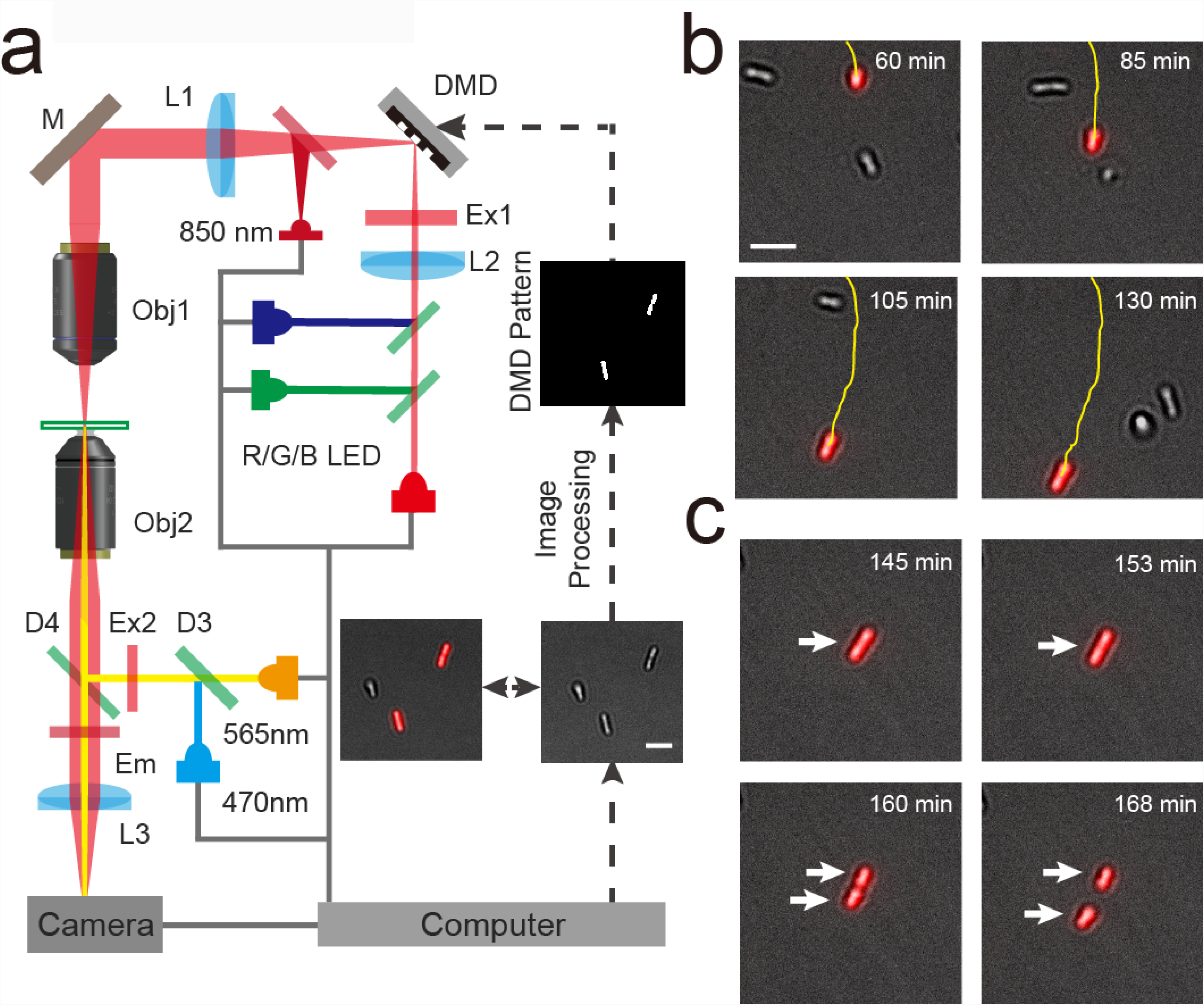
Using Adaptive Tracking Illumination Microscopy (ATIM) to exactly illuminate single *P. aeruginosa* cells on surface. (a) Schematic drawing of the ATI system. A high-throughput bacterial tracking algorithm was employed for analyzing cells’ behavior in real time and the information was immediately fed back to an adaptive microscope equipped with a digital micromirror device (DMD). (b) Example depicting one cell of interest being tracked and projected in real time. (c) The feedback illumination can generate projected patterns to exactly follow the daughter cells after the tracked cell divides. Scale bar for all images is 5µm.

**Figure 5 – Figure supplement 2.**
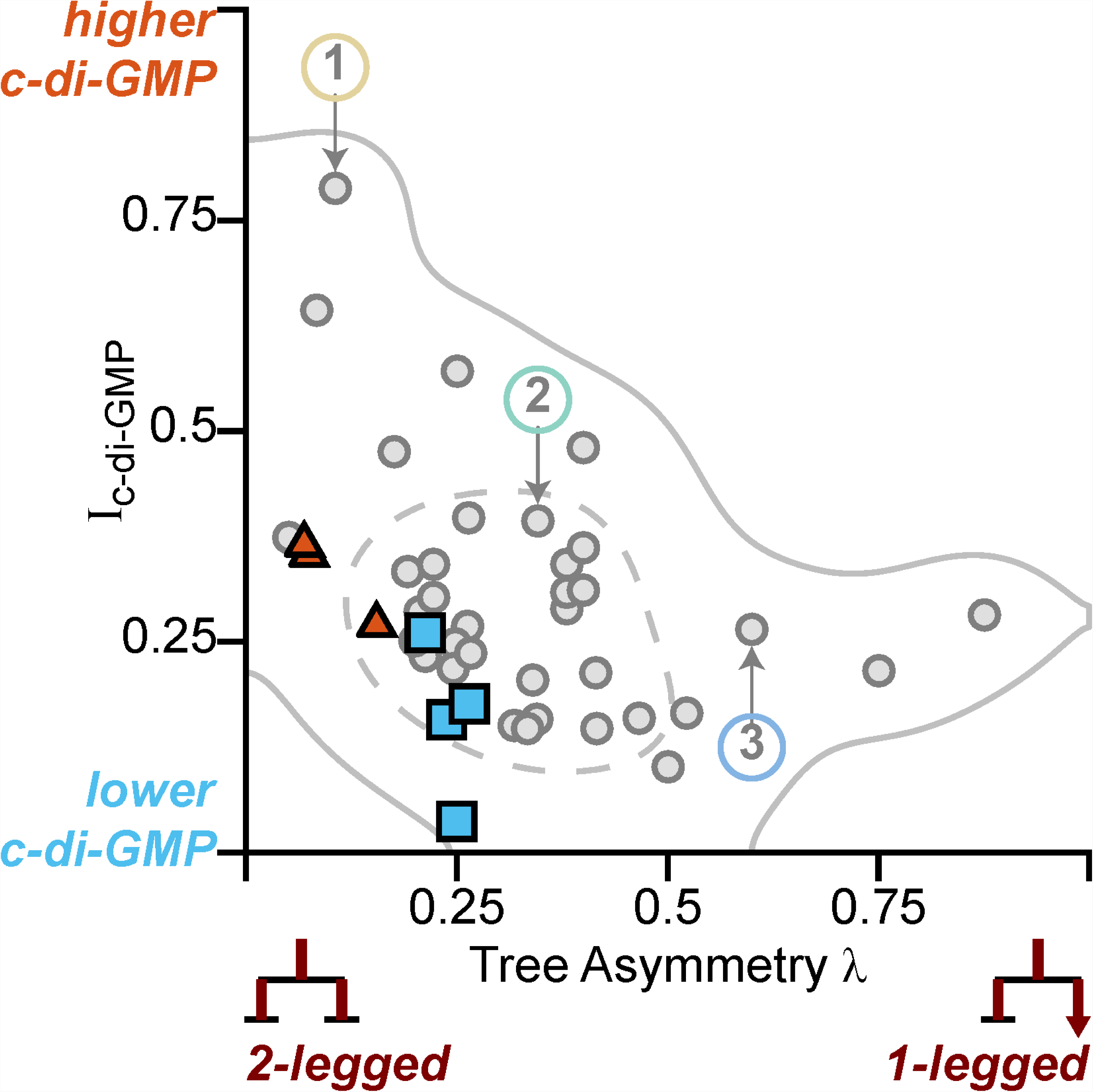
Optogenetic-controlled families follow the trend of family behavior observed in wt PAO1 cells, with illuminated families resembling the high c-di-GMP matrix producers and control families resembling low c-di-GMP surface explorers. Families plotted in Figure 5 (plus one additional control family) are plotted on top of Figure 4C (I_c-di-GMP_ vs λ). Red triangles and blue triangles represent the illuminated and control families, respectively, while the original data from Figure 4C is greyed out. Illuminated families have higher I_c-di-GMP_ and lower λ than the control families.

**Movie 1. Single cells are precisely illuminated by ATIM via *in situ* analysis and tracking of bacteria.** The left panel shows the merged images of *gfp*_ASV_ and mCherry fluorescence microscopy images over time. The right panel shows the merged images of red LED projected patterns and bright field images corresponding to the left panel. The fluorescence intensity of *gfp*_ASV_ in the illuminated cells and their offspring (colored red in right panel) is significantly increased after using ATI for 460 mins. In contrast, the *gfp*_ASV_ fluorescence intensity of the un-illuminated cells remains low and these cells remain motile.

